# Mapping the mutational landscape of a full viral proteome reveals distinct profiles of mutation tolerability

**DOI:** 10.1101/2024.03.07.583990

**Authors:** Beatriz Álvarez-Rodríguez, Sebastian Velandia-Álvarez, Christina Toft, Ron Geller

## Abstract

RNA viruses have notoriously high mutation rates due to error-prone replication by their RNA polymerase. However, natural selection concentrates variability in a few key viral proteins. To test whether this stems from different mutation tolerance profiles among viral proteins, we measured the effect of >40,000 non-synonymous mutations across the full proteome of coxsackievirus B3 as well as >97% of all codon deletions in the non-structural proteins. We find significant variation in mutational tolerance within and between individual viral proteins, which correlated with both general and protein-specific structural and functional attributes. Further, mutational fitness effects remained largely constant across cell lines, highlighting conserved selection pressures. In addition to providing a rich dataset for understanding virus biology and evolution, our results illustrate that incorporation of mutational tolerance data into druggable pocket discovery can aid in selecting targets with high barriers to drug resistance.

## INTRODUCTION

All RNA viruses must execute a conserved and coordinated set of functions to successfully propagate, including the delivery, replication, and packaging of their genomic material inside host cells as well as the modification of the host environment to favor viral spread. To accomplish these tasks, viruses must encode a minimal repertoire of functionally conserved proteins, including one or more structural proteins involved in the formation of the viral particle (e.g. capsid or envelope proteins) and a non-structural protein required to replicate the genome, the RNA-dependent RNA polymerase^1^. In addition, most harbor additional non-structural proteins that are involved in various aspects of genome replication (e.g. helicases) and modification of the host environment (e.g. viral proteases, viroporins, etc.).

RNA viruses have a remarkable ability to rapidly evolve and adapt to changing environments. This is mostly fueled by extreme mutation rates stemming from replication by error-prone polymerases that lack proof-reading functions^2–4^. During replication, these polymerases frequently misincorporate nucleotides, resulting in amino acid (AA) substitutions, but can also introduce insertions and deletions via distinct mechanisms^5^. Experimental assessment of viral polymerase mutation rates in the absence of selection pressures for viral fitness has shown that mutation rates remain high across the majority of the genome^6^. However, in nature, where selection purges deleterious mutations, mutational diversity is not uniformly distributed across different viral protein classes. Specifically, viral proteins targeted by adaptive immune responses, such as viral capsids or envelope proteins, can show greater sequence diversity than other protein classes^7–9^. While previous studies have shed light on the overall distribution of mutational fitness effects (MFE) between structural and non-structural proteins^10^, a comprehensive analysis of how different viral protein classes accommodate mutations remains lacking. Furthermore, our knowledge of how deletions affect viral protein function has only recently begun to be addressed and remains limited^5,11–13^.

Within this framework, Coxsackievirus B3 (CVB3) provides an excellent model for studying mutational tolerance across different viral protein classes. CVB3 is a human enterovirus belonging to the picornavirus family. Picornaviruses are a ubiquitous family of positive-strand RNA viruses infecting animals and humans, imposing significant economic and health burdens^14^. The CVB3 genome encodes an array of structurally and functionally diverse proteins, including four structural proteins (VP1-4) that form the viral capsid and seven non-structural proteins (2A-2C and 3A-3D) that play diverse and essential roles in viral replication and host modulation. The structure and function of most CVB3 proteins have been extensively studied and high-resolution structures of CVB3 or related enterovirus proteins are available. Importantly, many CVB3 proteins show structural conservation across the greater picornavirus family and even other viral families (e.g. proteases or polymerases), facilitating the extrapolation of results to additional pathogens.

Deep mutational scanning (DMS) approaches provide comprehensive insights into MFE by systematically introducing a large fraction of all single AA mutations into a gene of interest and measuring their effects on fitness^15^. Previously, we applied DMS to the CVB3 capsid region to define MFE and their correlation with structural, functional, and evolutionary parameters^16^. Herein, we extend our DMS analysis to the non-structural proteins (~60% of the proteome). Together, we assess the effect of >40,000 AA mutations across the full viral proteome as well as >97% of codon deletions across the non-structural proteins. Our results provide a comprehensive full-proteome dataset of MFE across two cell lines, unveiling both general and protein-specific responses to mutations and deletions. In addition, we show that MFE data can be integrated into drug discovery to identify druggable pockets with high barriers to the development of drug resistance.

## RESULTS

### Deep mutational scanning of the CVB3 proteome

Previously, we used a PCR-based mutagenesis protocol to introduce 92% of all possible AA mutations across the capsid of CVB3 and analyzed their effects on viral fitness^16^ (Figure 1A). To understand how different protein classes accommodate mutations, we extended our analysis to the remainder of the proteome, including all non-structural proteins. For this, DMS was performed on the P2 and P3 regions of the proteome independently, with short overlaps in the downstream (P1, 5 AA) and upstream (P3, 20 AA) regions included in the P2 mutagenesis region to enable standardization across all three regions. Rather than using the previously employed PCR-based mutagenesis method, a new synthetic biology approach was utilized for the mutagenesis, where ~250bp-long oligonucleotides encoding either the deletion of a given codon or a single mutant codon for each of the 19 possible non-synonymous mutations are cloned into the infectious clone (Figure 1B). In addition, stop codons and synonymous codons were introduced as controls at low frequencies. As compared to PCR-based mutagenesis, this method should significantly reduce redundancy (n = 19 versus n = 64 mutant codons with random mutagenesis), minimize background from non-mutagenized WT sequence, and limit the occurrence of multiple mutations per clone (see Figure S1). Importantly, it enables the selection of mutant codons that are the greatest number of base changes away from the original codon, overcoming background from sequencing errors that are dominated by single substitutions^16^.

**Figure 1.**
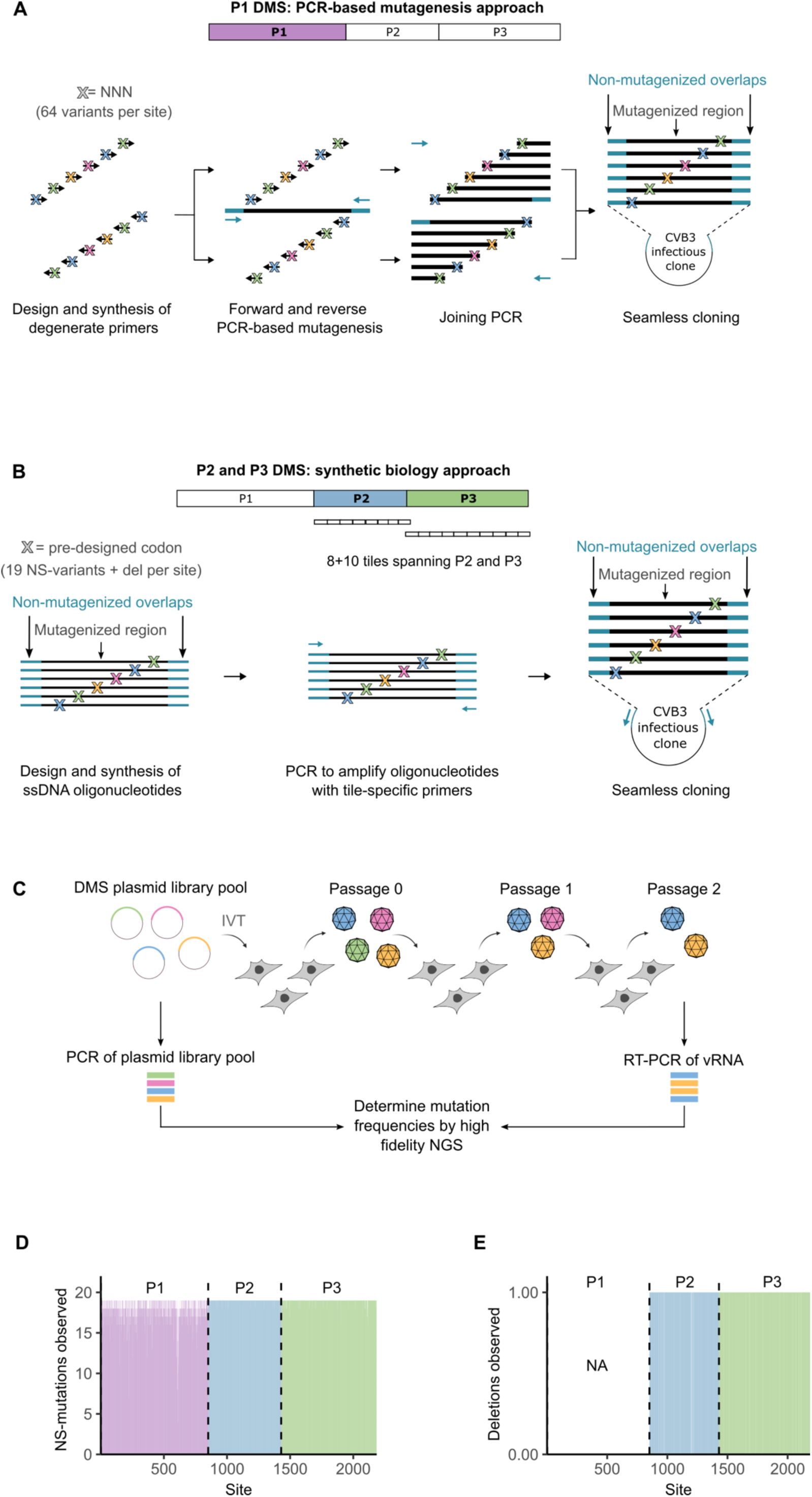
Overview of the full-proteome DMS analysis. **(A)** Overview of the mutagenesis protocol used for the structural proteins. A codon-level PCR mutagenesis using degenerate primers with randomized bases at the codon-matching positions (NNN) was used to mutagenize the complete P1 region, as previously reported^16^. **(B)** Overview of the mutagenesis protocol used for the non-structural proteins. The P2 and P3 regions were mutagenized separately via a synthetic biology approach. Each region was divided into ~250bp tiles. For each residue to mutagenize in the tile, a pool of oligonucleotides was synthesized encoding either a deletion or a single mutant codon for each of the possible 19 AA mutations. Stop codons and silent mutations were introduced as controls at a low frequency. All oligonucleotides harbored complementary sequences to the flanking regions, which were used to both amplify the tiles from the pool and clone them into the corresponding region in the infectious clone. **(C)** Generation of viral populations and determination of MFE by next-generation sequencing (NGS). Viral RNA (vRNA) was produced from the mutagenized plasmid of each region by in vitro transcription (IVT) and electroporated into HeLa-H1 cells. Following a single cycle, cells were freeze-thawed to liberate viral particles. Two additional rounds of infection were performed at low multiplicity of infection to reduce complementation and enable selection. The mutagenized region was then amplified from both the plasmid libraries as well as from viral RNA from passage 2 following reverse transcription, and mutation frequencies were obtained via high-fidelity NGS. **(D)** The number of AA mutations observed for each residue. **(E)** The number of single codon deletions observed for each residue across the non-structural region (P2 and P3). Note, deletions were not included in the mutagenesis of the P1 region. See GitHub section A2 for sequences of synthetic oligonucleotides, Table S1 for primers used to amplify by PCR the vector and the tiles and Table S2 for NGS statistics.

Three independent mutagenized libraries were generated for the P2 and P3 regions. These were used to produce viral RNA by in vitro transcription, which was electroporated into HeLa-H1 cells to generate viral populations. Two subsequent rounds of infection (passages) were then performed at a low multiplicity of infection to reduce complementation and enable selection for fitness (Figure 1C). For the P1 region, previously characterized passage 1 viral populations^16^ were used to similarly infect cells for a second round. Mutation frequencies were then obtained from both the mutagenized libraries and the passage 2 viral population using a high-fidelity next- generation sequencing (NGS) technique^16^. From this, we derived the mutational fitness effects (MFE) of 97.6% (42,630) of all AA mutations across the full proteome and 97.8% (1,305) of all single deletions across the non-structural proteins (Figure 1D, E). Sequencing statistics are available in the supplementary Table S2.

To calculate MFE, the relative frequency of each AA mutation versus the WT AA at each site in the viral populations was divided by its relative frequency in the plasmid libraries and averaged across the independent replicates (Figure 2A; See Figure S2 for correlations between replicates). Examination of MFE for mutations present in the overlap regions between P1/P2 (n = 61) and P2/P3 (n = 161) showed a good linear relationship (Pearson correlation coefficient = 0.74 and 0.83 for P1/P2 and P2/P3, respectively; Figure 2A and S3). This allowed us to normalize MFE across the three regions based on a linear model. Finally, to validate our results and calibrate selection in our experiments to a more realistic measure of fitness, we generated 31 individual mutants across the full proteome and experimentally determined their fitness by direct competition with the WT virus. DMS-derived MFE correlated well with experimental measurements of fitness (Pearson correlation coefficient = 0.78, p< 10^−7^; Figure 2A), enabling us to standardize our DMS results to experimentally determined fitness values using a linear model (see Methods).

**Figure 2.**
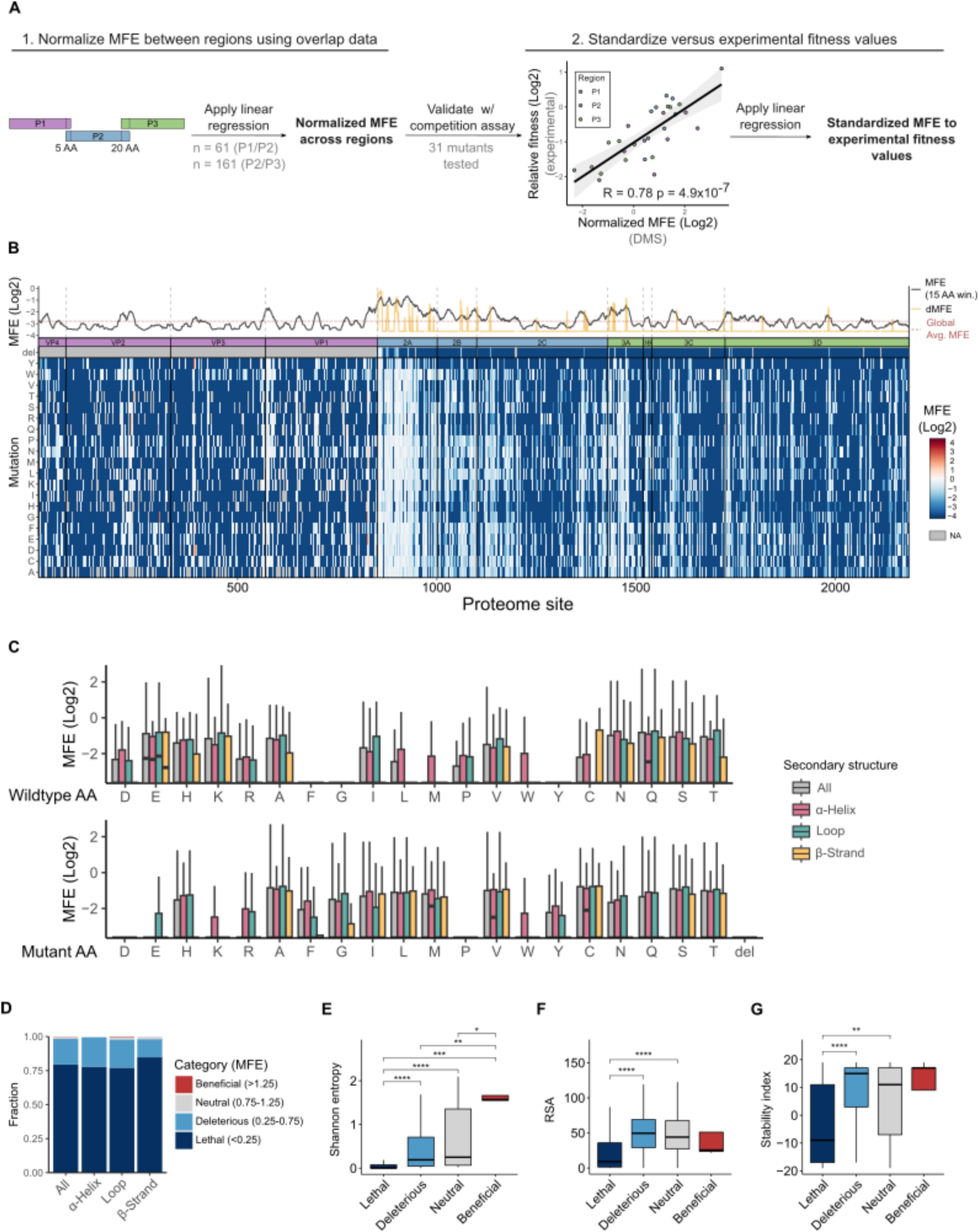
Overall patterns of MFE across all the CVB3 proteome. **(A)** Overview of the MFE normalization protocol between the three mutagenized regions and their calibration to experimentally measured fitness values. A five and 20 AA overlap with the P1 and P3 region, respectively, was included in the mutagenized P2 region. Linear models based on the MFE of mutations in these overlapping regions were used to normalize the P1 and P3 MFE relative to those of P2. These overlap-normalized MFE were then calibrated to experimentally-measured fitness values of 31 individual mutants across the proteome using a linear model, as these showed high correlation (Pearson correlation coefficient = 0.78, p < 4.9×10^−7^). **(B)** Proteome-wide MFE distribution. Top panel: line plot showing a 15 AA sliding window analysis of MFE (black line), individual deletion MFE (dMFE; yellow line), or the global average MFE across the full CVB3 proteome (dashed red line). Bottom: Heatmap of MFE and dMFE across the full CVB3 proteome. **(C)** The distribution of MFE based on the nature of the WT AA or the mutation relative to different secondary structure elements. **(D)** The frequency of lethal, deleterious, neutral, or beneficial mutations across the full proteome (all) and according to secondary structure element. **(E-G)** The distribution of sequence variability in enterovirus B sequences (Shannon entropy) **(E)**, relative surface area (RSA) **(F)**, or stability index **(G)** of each residue relative to different fitness categories. ns: p > 0.05, *p < 0.05, **p < 0.01, ***p < 0.001, ****p < 0.0001 by Mann-Whitney test following multiple test correction. See Tables S3 and S4 for mutation and site data, respectively. See GitHub section B4 for calculation and correction of MFE and section A4 for experimental data of the competition assay.

### Overall patterns of MFE across the CVB3 proteome

MFE varied across and between individual proteins, with clear regions of tolerance to mutations (e.g. N-terminal region of VP1, 2A, and 3A; Figure 2B and Tables S3, and S4). Overall, the nature of the WT AA and the introduced mutation had a strong impact on MFE, which was further influenced by protein secondary structure for some AA. Specifically, mutations at five AA (M, F, G, W, and Y) had a strong fitness cost independent of the secondary structure, as were mutations from any AA to D, P, or a deletion (Figure 2C). The remaining AA residues showed differential MFE depending on secondary structure elements (e.g. mutation at L, M, and W at α-helices or mutation to E in loops; Figure 2C). Interestingly, the MFE of certain AAs varied depending on whether the residue was undergoing mutation or if the mutation involved the AA itself. For example, F, G, and Y exhibited intolerance to mutations but were surprisingly tolerated as mutations themselves, reflecting their strict functional and structural roles as WT residues (Figure 2C). On the other hand, the AA D and P displayed the opposite effect, showing a higher tolerance to accommodate substitutions than to be introduced as mutations (Figure 2C).

To facilitate the comparison of our results with evolutionary and structural parameters, MFE were grouped into fitness categories as follows: lethal (MFE of 0–0.25), deleterious (MFE of 0.25–0.75), neutral (MFE of 0.75–1.25), and beneficial (MFE >1.25; Figure 2D). Comparison of fitness categories with natural sequence variation (Shannon entropy) in enterovirus B sequences revealed lethal mutations to concentrate in invariable residues while the opposite was observed for beneficial mutations, highlighting general agreement between lab-measured MFE and natural selection processes (Figure 2E). In line with this observation, the incorporation of MFE into phylogenetic models outperformed standard models (YNGKP/Goldman-Yang models)^17^ for all three regions (Table S5), revealing that DMS-derived MFE reflect selection processes in nature. As previously observed for the capsid region^16^, multiple additional structural attributes were also correlated with MFE. Specifically, MFE differed by protein secondary structure, with loops having the largest fraction of neutral and beneficial mutations while β-strands had the largest fraction of lethal mutations (p < 10^−15^ by Fisher’s exact test for all; Figure 2D). In addition, lethal and deleterious mutations were enriched in buried residues (relative surface area, RSA, <25; Figure 2F) and had a larger fraction of destabilizing mutations (Figure 2G).

### MFE distribution within and between different protein classes

A general reduction in fitness was observed upon mutation across all CVB3 proteins (Figure 3A). However, clear differences in the ability to tolerate mutations were observed between individual proteins (p<10^−16^ by Kruskal-Wallis test for all; Figure 3A). The icosahedral CVB3 capsid is the most complex of all CVB3 protein structures. It is formed by the stepwise assembly of 60 copies of each of the four capsid subunits around the viral genome, with VP1-3 forming the external surface and VP4 lining the inner surface^18^. Of all CVB3 proteins, the structural proteins (VP1-VP4) had the largest fraction of lethal mutations but were the only ones harboring beneficial mutations (Figure 3B). Overall, surface-exposed residues in the external capsid proteins (VP1- VP3) were more tolerant to mutations than buried residues (Figure 3C, Figure S4A), with beneficial mutations (n = 230 mutations in 97 sites) enriched in surface-exposed loops (88.7%; p < 10^−9^ by Fisher’s test). The only sites in the proteome where mutations were on average beneficial (average site MFE > 1.25; n = 5) were found in the external capsid proteins. These sites belong to known antibody neutralization epitopes^19^, likely reflecting the absence of this selective pressure in cell culture and highlighting its relevance to evolution in nature.

**Figure 3.**
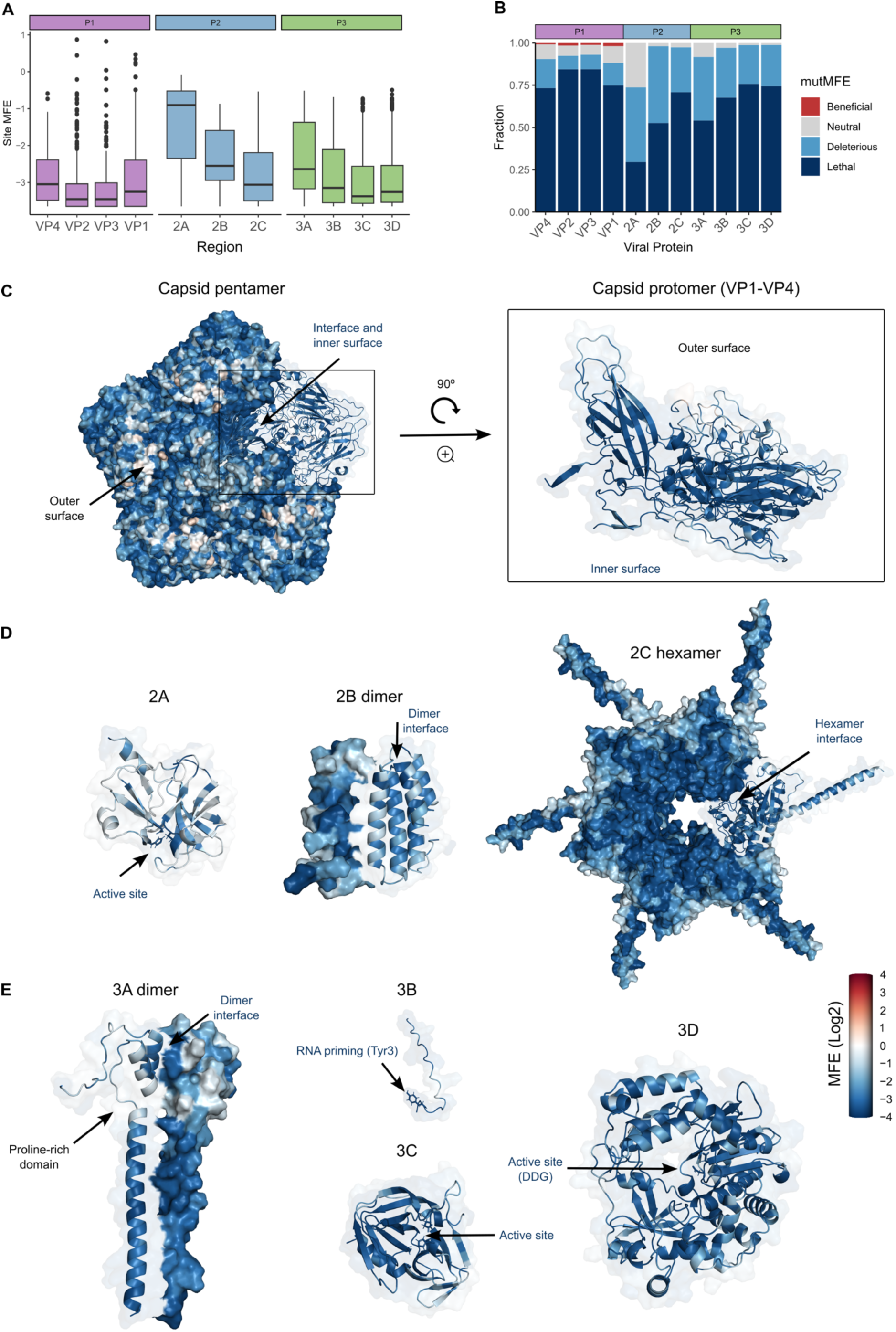
MFE distribution across the full CVB3 proteome. **(A)** Boxplot of the average site MFE across all CVB3 proteins. **(B)** The distribution of fitness categories across all CVB3 proteins. **(C-E)** The CVB3 capsid pentamer **(C)**, P2-derived proteins **(D)**, or P3-derived proteins **(E)** colored according to the average MFE of each site. All P2 and P3 structures were predicted using Alphafold2 according to their known quaternary structure. For 2B, a dimer is shown as the tetramer prediction did not reflect the expected structure. See GitHub section A7 for Pymol sessions of MFE mapped onto protein structures.

Viral proteases are responsible for the maturation of viral proteins from larger precursors and/or the cleavage of cellular factors to facilitate viral growth. CVB3 encodes both a chymotrypsin-like (2A) and a trypsin-like (3C) protease^20,21^. The 2A protease is only required for the cleavage of the capsid precursor P1^21,22^, while 3C mediates the remaining nine cleavage events at conserved Q-G/N residues^20^. Both proteases also cleave a non-overlapping array of cellular proteins to favor viral replication^20–23^. Comparison of MFE between these proteases revealed opposite mutation tolerance profiles, with 2A being the most, and 3C among the least, tolerant to mutations of all proteins (Figure 3A, B, D, E). In both proteases, mutations in the active site were highly deleterious (Figure S4C). However, a single mutation at each catalytic triad residue in 2A (H21Q, D39Q, C110W) or 3C (H40Q, E71D, C147W) was still observed in the viral populations at low frequency (maximal fitness of 0.19 and 0.32 for 2A and 3C, respectively; Table S4), suggesting some tolerance to mutations at these critical sites. As expected, mutations at the conserved Q residue of 3C cleavage sites were also highly intolerant to mutations (Figure S4D). However, Q to H substitution at these sites was also observed at low frequency in the viral populations (MFE < 0.25 reduction versus WT) in the non-structural proteins but not in the capsid. This difference could be attributed to the nature of the protease mediating the cleavage, as the structural proteins are cleaved only by the 3CD precursor rather than the mature 3C protease^24^. Alternatively, this could be related to the more complex maturation process of the capsid proteins, which requires the interaction with cellular protein folding factors that can dictate the fitness landscape of the capsid^25^.

All positive-strand RNA viruses replicate on viral membranes. To facilitate their replication on membranes, RNA viruses induce profound rearrangements of cellular compartments^26^. In CVB3, the 2B, 2C, and 3A proteins have all been shown to alter cellular membranes by distinct mechanisms^27–30^, although each protein also harbors additional activities. The CVB3 2B protein belongs to the viroporin family of oligomeric proteins complexes^30^. Viroporins regulate ion permeability by inserting into membranes, where they form ion channels, and play multiple roles in viral replication and pathogenesis^30,31^. The 99 AA long CVB3 2B protein has an amphipathic α-helix connected to a second α-helix by a short loop^32^ (see Figure 3D for the dimeric structure). Overall, 2B showed a low tolerance to mutation, with few neutral mutations (2%) and a large faction of deleterious (45.4%) and lethal (52.6%) mutations. The connecting loop was significantly more tolerant to mutation than both α-helices (p<10^−6^ by Mann-Whitney test; Figure S4G).

The viral protein 2C, or its precursor 2BC, can induce membrane rearrangements when expressed in cells^27^. However, 2C plays numerous additional functions in viral replication, including uncoating, RNA replication, genome encapsidation, and morphogenesis^33^. The structure of 2C contains an N-terminal membrane-binding domain (MBD), a central ATPase domain of the Superfamily 3 AAA+ helicase family, a cysteine-rich domain (zinc finger domain), and a C-terminal helical domain. High-resolution structures have shown 2C to form a hexameric ring complex^34^, which is essential for its ATPase activity. The different 2C domains showed strong variation in tolerance to mutations, with the MBD being the most accommodating of mutations and the ATPase domain the least (Figure 3D and Figure S4I). As expected for a multimeric protein, interface residues showed increased sensitivity to mutation (p<10^−16^ by Mann-Whitney test; Figure S4E). The 2C region also encodes an RNA structure that is essential for genome replication, the cis-regulatory element (CRE). Mutations in this region incurred a higher cost to mutation than the remainder of the protein (p<10^−15^ by Mann-Whitney test versus the rest of 2C; Figure S4F), likely reflecting disruptions to this important RNA structure.

The CVB3 3A protein, on the other hand, disrupts membranes by blocking anterograde transport between the endoplasmic reticulum (ER)-Golgi intermediate compartment (ERGIC) and the Golgi by preventing the formation of coatomer protein complex I (COPI)-coated transport vesicles^28^. It is a dimeric protein with an N-terminal domain (NTD) that is important for disrupting protein trafficking, a membrane binding domain (MBD), and a C-terminal domain (CTD). The 3A protein was the most tolerant to mutations following the 2A protease, with the NTD showing significantly higher cost to mutations relative to the MBD and CTD (p<10^−16^ by Mann-Whitney test; Figure 3E and S4H). In agreement with this observation, mutations in the NTD that allow for viral replication without blocking membrane trafficking have been reported^29,35^.

A defining feature of RNA viruses is the RNA-dependent-RNA polymerase (RdRP), which provides the essential function of replicating RNA genomes without utilizing a DNA intermediate. Overall, these proteins show a common fold that is even conserved in viral and non-viral RNA and DNA polymerases^36,37^. This includes a catalytic palm subdomain harboring the catalytic GDD residues, finger subdomains that bind RNA, and the thumb subdomain that interacts with the dsRNA products. In the CVB3 RdRP (3D), the index finger showed the highest tolerance to mutation compared to all other subdomains (Figure 3D and Figure S4J). Interestingly, the index domain is also more variable in nature than the other subdomains, except for the thumb subdomain (Table S3). As expected, all mutations in the catalytic GDD residues were invariably lethal, except for conservative substitutions of D329E and D330E, which nevertheless resulted in a drastic reduction of fitness (>70% reduction vs WT). Finally, several positive-strand RNA viruses encode a small protein that is covalently attached to the viral genome (viral protein genome-linked; VPg) and is used to initiate replication by the RdRP, including picornaviruses, potyviruses, and caliciviruses. A tyrosine residue in these VPgs becomes uridylated, which provides a free hydroxyl that can be extended by the viral polymerase, enabling a primer-free replication mechanism^14^. As expected, all mutations at this critical residue in the CVB3 VPg (3B protein) were lethal (Figure 3E).

### Analysis of single codon deletions across all non-structural proteins

We next analyzed the effect of single codon deletions on viral fitness (deletion MFE; dMFE) across the non-structural proteins. Overall, deletions were significantly more deleterious than mutations across both the P2 and P3 regions, with 96% of deletions reducing fitness below 25% of WT (Figure 4A, B). However, all non-structural proteins harbored sites where deletions were not lethal (dMFE > 0.25; Figure 4B). As with MFE, the most tolerant region was found in 2A, with 27 sites maintaining viability upon deletion (Figure S5A). In addition, an 8 AA stretch at the N terminus of 2A (residues 2-10) retained an average fitness of 81.8% (range, 68.4%-100%; Figure S5B). This likely reflects flexibility in the 2A cleavage site sequence requirements. In agreement with this, three of the five residues in the C terminus of VP1 that were included as part of the overlap with the P2 mutagenesis region also tolerated deletions (positions −5, −3, and −2 relative to AA 1 of P2; Table S4). Additional residues with viable deletions were found in 2C and 3A (n = 8 each), 2B (n = 4), and 3C and 3D (n = 2 each; Figure S5B, C). Interestingly, nearly 4% of deletions (51/1,305) were better tolerated than certain mutations at the same residue (dMFE > minimum MFE [minMFE]; Figure 4C). Hence, the structural and biochemical properties of some AA can be more deleterious than the absence of a residue, as previously observed in a smaller study^12^. Such sites were significantly more variable in nature (Figure 4D) and had specific structural attributes: high relative surface exposure (Figure 4E) and probability of stabilizing protein structure upon mutation (Figure 4F) as well as an enrichment in loops and depletion from α-helices (Figure 4G). The AA D and P were particularly enriched at such sites (>2.5-fold versus sites with dMFE < minMFE), where specific mutations were more deleterious than deletions (e.g. D to K, or P to F).

**Figure 4.**
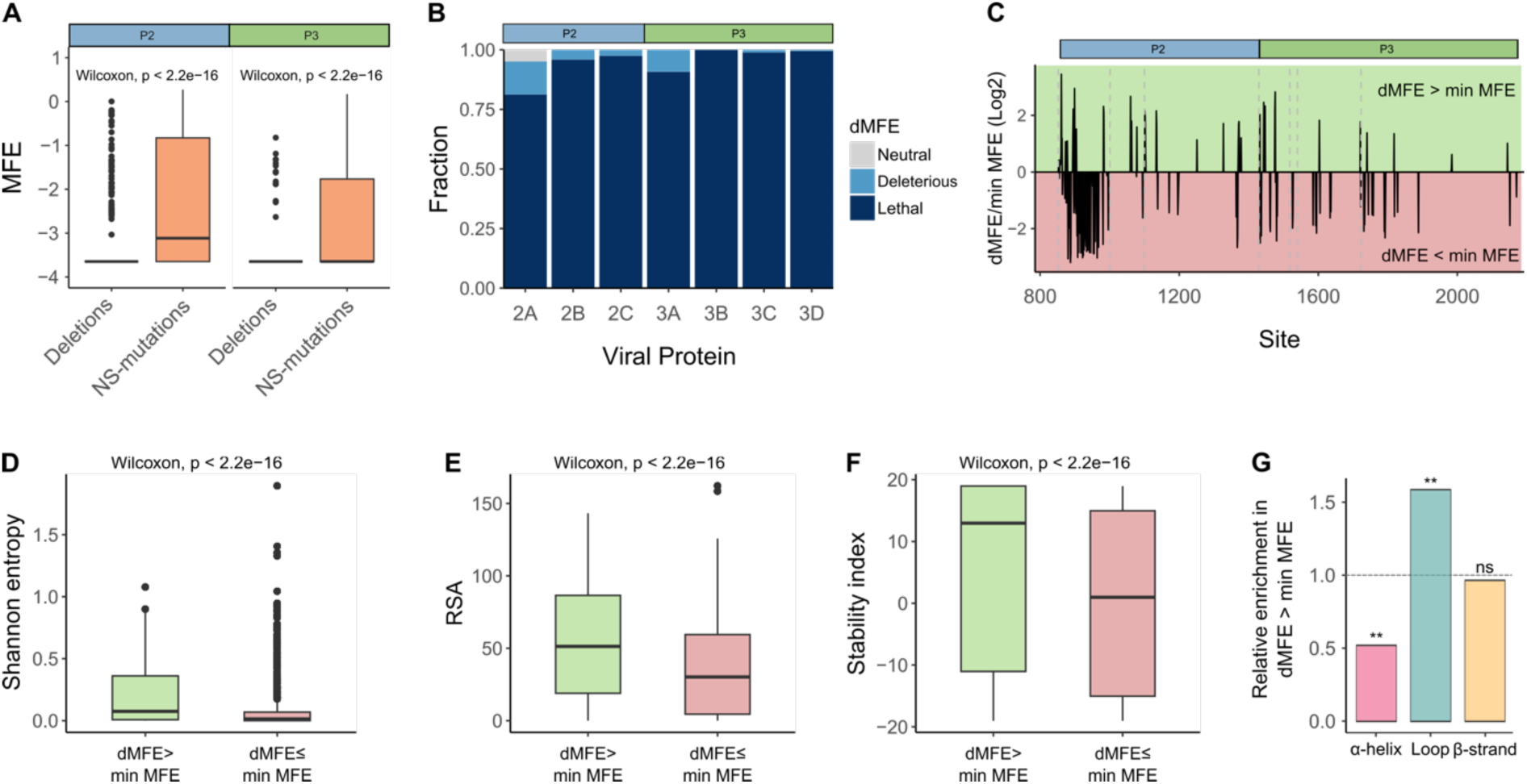
Analysis of single codon deletions across all non-structural proteins. **(A)** The effects of non-synonymous mutations (NS-mutations) versus deletions on viral fitness across the P2 and P3 regions. **(B)** The distribution of deletions according to their fitness categories across all non-structural proteins. **(C)** Deletions can be better tolerated than mutations. The ratio of dMFE to the most detrimental MFE at each site (minMFE), with positive values indicating residues in which deletions were better tolerated than some mutations. **(D-G)** Deletions that are better tolerated than mutations (dMFE > minMFE) occur at sites that are more variable in nature **(D)**, more surface exposed **(E)**, have a larger fraction of mutations that are stabilizing **(F)**, and show differential distribution at secondary structure elements **(G)**. ns p > 0.05, **p < 0.01.

### MFE remain constant across cellular environments

During colonization of the host, viruses must frequently infect different tissue environments, which can present distinct selection environments^38^. To better understand whether MFE observed in HeLa-H1 cells are generalizable across different cellular environments, we obtained MFE across the full CVB3 proteome in an additional cell line, hTERT-immortalized retinal pigment epithelial cells (RPE). These were chosen as they provide a significantly more restrictive environment for CVB3 than HeLa-H1 cells (116 ± 12.9-fold infectivity reduction) and mount a stronger type I interferon response (maximal inhibition of virus infection of 82.1±6.6% and 21.8±6.3% for RPE and HeLa-H1 with 20 IU of interferon-β, respectively). Hence, RPE cells could present a distinct environment for CVB3 than HeLa-H1 cells.

To identify sites showing different fitness profiles across the two cell lines, RPE cells were infected with the mutagenized passage 1 virus populations from all three regions (P1, P2, and P3) produced in HeLa-H1 cells to obtain RPE-derived passage 2 virus populations (Figure 5A). Mutation frequencies in these populations were then obtained using high-fidelity NGS as before and compared to those obtained in HeLa-H1 cells (See Table S6 for sequencing statistics and Figure S6 for correlations between replicates). For this, the relative frequency of each mutation compared to the WT AA at each site in viral populations derived from HeLa-H1 cells was divided by that observed in virus populations derived from RPE cells, yielding a differential selection score for each mutation (Table S7). Finally, all mutations in each site showing enhanced fitness in a HeLa-H1 cells (differential selection score >1) were summed to provide a site differential selection score that reflects the overall contribution of a site to improved fitness in this cell line (Figure 5B; Table S7). Similarly, mutations with improved fitness in RPE cells (differential selection score <1) were summed to reveal sites showing improved fitness in RPE cells (Figure 5B; Table S7). Overall, sites across the non-structural regions did not show strong differences in fitness between the two cell lines (Figure 5B), indicating similar selection pressures were present in both environments. In contrast, multiple residues across the capsid regions showed large fitness grains (high site differential scores) in HeLa-H1 (Figure 5B). Except for one mutation in the internal capsid protein, these sites localized to surface-exposed loops, suggesting a role in receptor binding. As CVB3 can utilize two receptors for cell entry, the coxsackievirus-adenovirus receptor (CAR) and decay-accelerating factor (DAF; CD55)^39,40^, we examined whether these sites colocalized with the binding footprint of either receptor. Indeed, most sites overlapped with DAF-binding residues (Figure 5B, C). Furthermore, DAF was readily detected by Western blot in HeLa-H1 cells but was undetectable in RPE cells, while CAR expression was similar between both cell lines (Figure 5D, Figure S7). Hence, cell-specific entry factors impose strong selection pressures between different cell lines.

**Figure 5.**
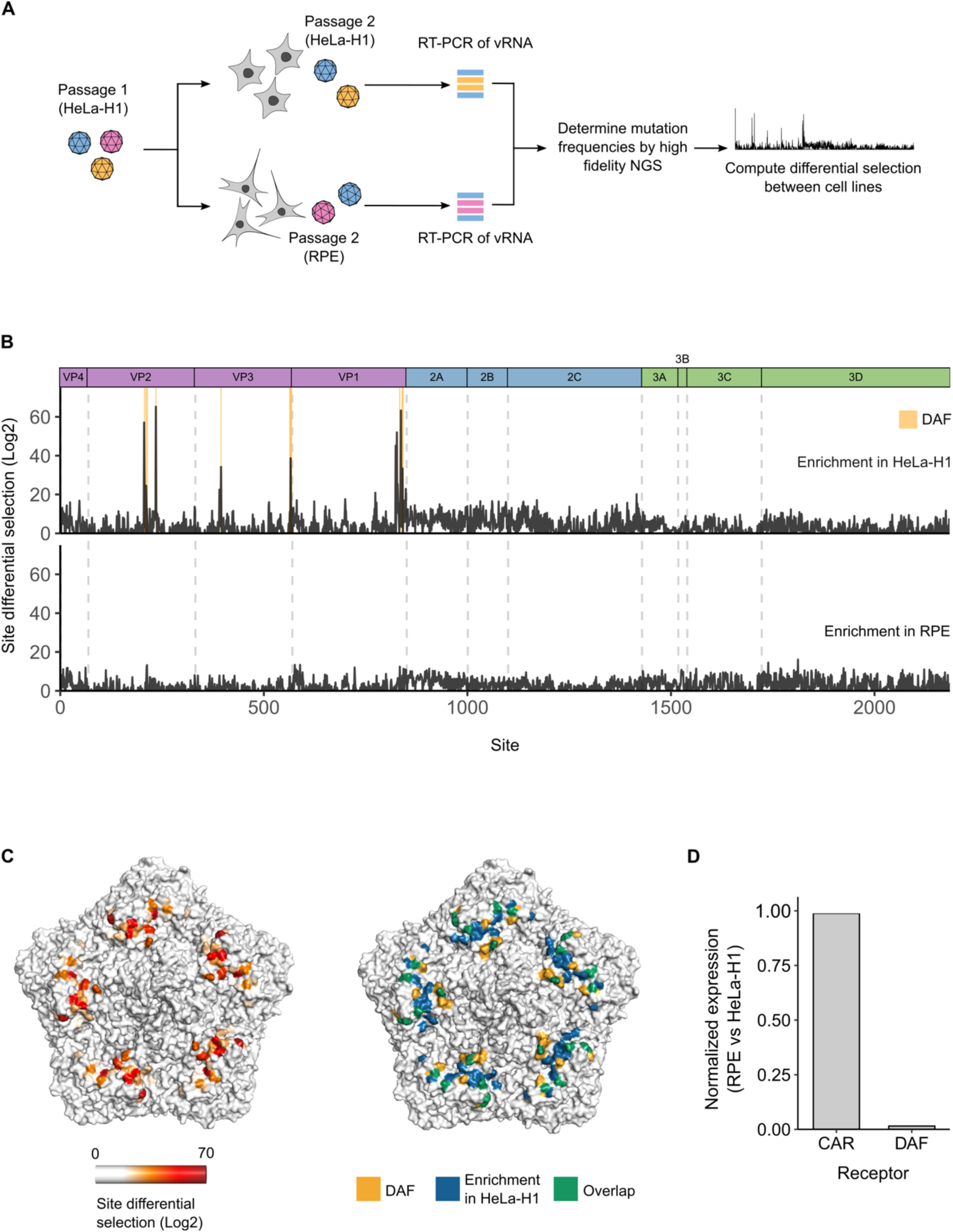
MFE are conserved across cell lines. **(A)** Overview of the experimental design for comparing MFE across HeLa-H1 and RPE cells. **(B)** Site differential selection for HeLa-H1 (top) and RPE cells (bottom). Residues showing large site differential selection scores that overlap with the DAF (CD55) footprint are highlighted in yellow. **(C)** The CVB3 capsid pentamer structure colored by the site differential selection score (left panel) or by sites showing large site differential selection scores in HeLa-H1, the DAF footprint, and their overlap (right panel). **(D)** Relative expression of DAF and CAR in RPE cells versus HeLa-H1 cells. Protein expression in each cell line was assessed by western blot and standardized to GAPDH expression. See Table S6 for NGS statistics and S7 for differential selection values between cell lines and Figure S7 for the western blots.

### Identification of mutation-intolerant druggable pockets

The extreme mutation rates and large population sizes of RNA viruses generate tremendous diversity during infection. As a result, antiviral therapies can be rendered inefficacious by the selection of variants encoding drug-resistant mutations that maintain sufficient replicative fitness to spread in the host. A potential means of preventing the emergence of drug resistance is the targeting of druggable pockets that accommodate few viable mutations. To evaluate if different druggable pockets show distinct mutation tolerance profiles, we combined computational druggable pocket predictions with our MFE information across the full CVB3 proteome (Figure 6A; Table S8). In total, 12 druggable pockets were predicted, which were distributed across the capsid (n = 5), 2A (n = 1), 2B (n = 3), 2C (n = 1), and 3D (n = 2; Table S8) proteins. Mapping of MFE to these druggable pockets revealed strong variation in mutational tolerance (Figure 6B; Table S8). Specifically, drug pocket 12 in the 3D polymerase was highly intolerant to mutation, having only a single neutral mutation out of 679 possible mutations in its 34 residues, with the remaining mutations being either deleterious (14%) or lethal (86%; Figure 6B, C). In contrast, druggable pocket 6 in 2A had 54 neutral or beneficial mutations across 18 of its 24 residues (75%), suggesting a lower barrier to the development of drug resistance (Figure 6B, C). Similarly, all druggable pockets in the capsid had a significant fraction of neutral mutation (3.1% – 7.9%), with 80% of them even having beneficial mutation (0.4% –1.2% of all mutations; Figure 6B). Hence, the incorporation of MFE data into antiviral target selection could help identify drug targets with higher barriers to drug resistance.

**Figure 6.**
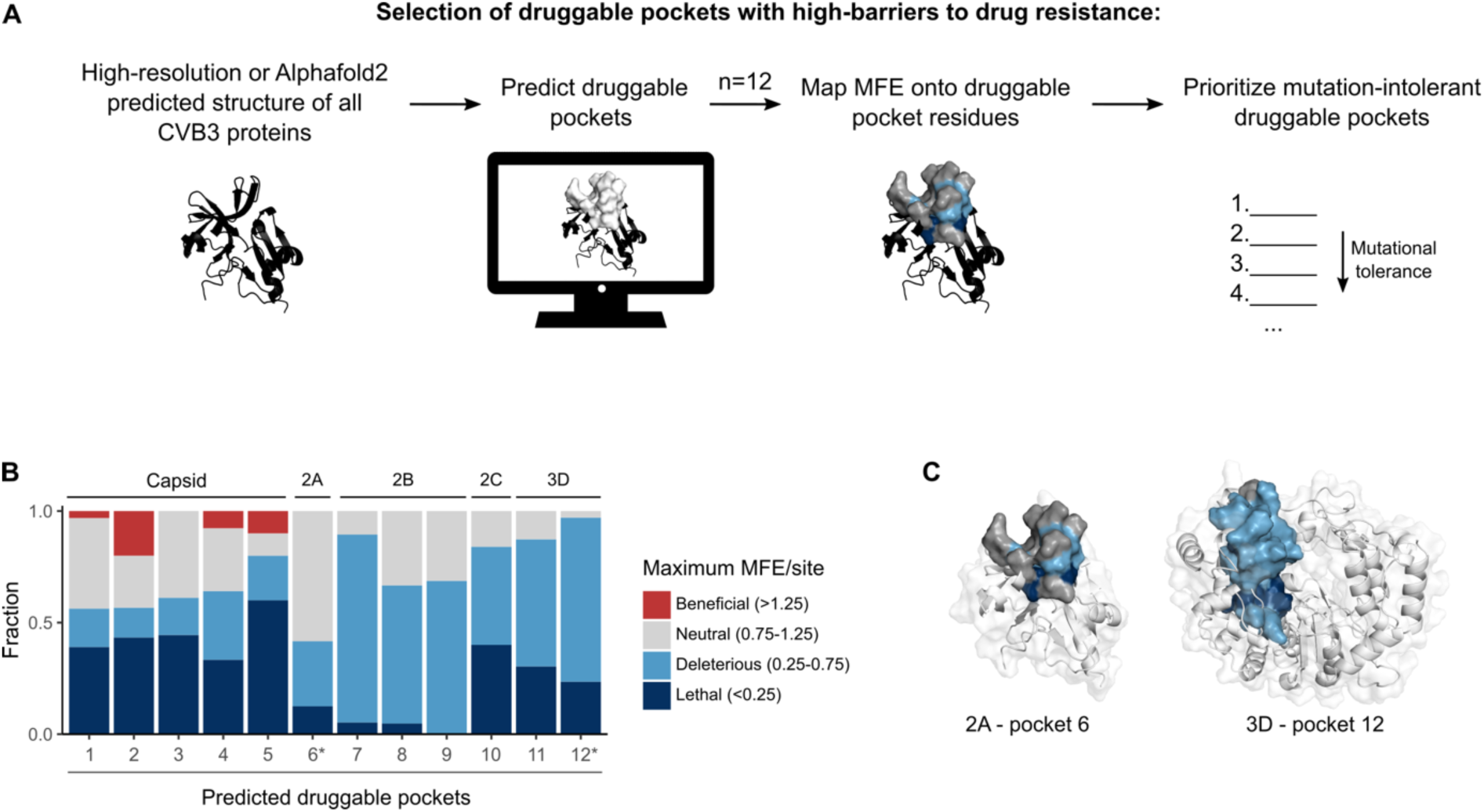
MFE can inform druggable pocket selection. **(A)** Overview of the pipeline for prioritization of druggable pockets based on MFE data. **(B)** The fraction of residues in each drug pocket belonging to a given fitness category based on the most permissive mutation at each residue (i.e. maximum MFE per site). Asterisks indicate drug pockets mapped onto the protein structure in (C). **(C)** Mapping of pocket 6 and 12 residues onto the 2A and 3D protein structure, respectively, and colored according to the fitness category of the most permissive mutation at each residue. See Table S8 for data on druggable pockets and MFE.

## DISCUSSION

The average rates of nucleotide misincorporation by the polymerases of RNA viruses have been experimentally determined and are generally orders of magnitude higher than other organisms^2–4^. These extreme mutation rates are believed to be the result of a tradeoff between replication kinetics and fidelity^41^, which RNA viruses have optimized to increase replication speed while avoiding population extinction due to mutational load. Aside from speed, high mutation rates confer RNA viruses the ability to rapidly evade immunity, develop drug resistance, and adapt to new hosts. As most mutations are deleterious, it has been suggested that viral proteins harbor unique properties to mitigate mutational fitness costs^42^. However, mutational diversity in nature is unevenly distributed across and within different viral proteins ^7–9^, suggesting distinct mutational tolerance across different viral protein classes. Indeed, RNA viruses encode an array of different protein classes, which vary in size (22 – 462 AA in CVB3) and complexity (1 – 240 subunits in CVB3), that could influence mutational tolerance.

A comprehensive analysis of how mutations affect fitness across different protein classes has not been reported. An analysis of how ~20% of all possible single AA mutations affected poliovirus fitness revealed differences in the overall distribution of MFE between structural and non-structural proteins, with the latter having a higher fraction of neutral and positive mutations^10^. However, this work largely focused on the effects of single mutation per codon, which limits the sampled mutational space, and did not analyze different proteins in detail. Rather than relying on the viral polymerase to generate mutation, deep mutational scanning (DMS) experimentally introduces a large fraction of all possible single AA mutations across proteins of interest. As such, it is ideal for obtaining a comprehensive understanding of MFE. Thus far, this technique has been used to define MFE in different individual proteins, including structural proteins from enveloped and non-enveloped viruses or non-structural proteins such as polymerase or matrix proteins^13,15^. However, a DMS analysis spanning a full proteome has yet to be reported. In this work, we utilized mutagenized viral populations from a previous DMS study of the CVB3 capsid region^16^ with a new mutagenesis protocol of the non-structural proteins to provide a comprehensive analysis of MFE across a full proteome. Importantly, we calibrated the DMS-derived MFE relative to experimental measurements of viral fitness to provide a more realistic measure of fitness that is lacking in the majority of published DMS studies, including our previous analysis of the CVB3 capsid.

Overall, we define the effect of >97% of all possible single AA mutations across the complete proteome of CVB3. Aside from providing a wealth of information with which to interpret the biology and evolution of CVB3 and related viruses, our results reveal both general and specific information on how different protein classes accommodate mutations. Firstly, a strong bias in MFE is observed depending on both the original and mutant AA, which was dependent on different secondary structure elements for certain AA (Figure 2C). Moreover, even for the same AA in the same secondary structure element, MFE could significantly vary depending on whether a given AA was being mutated from (i.e. the WT residue) or to (i.e. the mutation), highlighting strong structural and functional constraints (Figure 2C). As expected, external residues in the capsid and surface-exposed residues in the non-structural proteins were more tolerant of mutations, highlighting a general sensitivity of buried residues to mutation (Figure S4B). In terms of protein classes, the structural proteins showed a higher fraction of lethal mutants than the non-structural proteins but were also the only ones to have beneficial mutations (Figure 3A, B). The only sites showing an average positive effect of mutation on fitness were found at previously defined antibody neutralization sites, reflecting the strong selection of immune evasion on virus evolution. Of the non-structural proteins, the two proteases showed different MFE profiles, with 2A being the most tolerant to mutations of all proteins and 3C among the least (Figure 3A, B). While this difference could be due to relaxed selection pressures in cell culture, 2A is also more variable than 3C across enterovirus B sequences (p<10^−16^ by Mann-Whitney test; Table S3), supporting inherent structural and functional differences. As 2A only performs a single cut in the CVB3 proteome, as compared to nine performed by 3C, this protease could be less constrained. Indeed, the protease function of 2A has been replaced by a short ribosomal skip-sequence, or duplicated, in some picornavirus genomes^21^. The remaining non-structural proteins showed varying sensitivity to mutation, with particular regions and/or domains showing differential ability to accommodate mutations that were associated with protein-specific functions.

Unlike mutations, global evaluation of the effects of deletions on protein function has only recently begun to be studied^5,11–13^. Taking advantage of the new synthetic biology mutagenesis approach utilized in this work, we included all single AA deletions across the non-structural proteins along with all single mutations. In agreement with prior work^5^, we observed deletions to be more deleterious than mutations, with the vast majority being lethal (Figure 4A). However, some deletions were neutral in the 2A protein, which was also the most mutation-tolerant CVB3 protein (Figure 4C). These were concentrated at the N-terminus of 2A, where this protease cleaves the downstream capsid region, highlighting flexibility in the sequence requirements of this protease. Interestingly, nearly 4% of deletions had higher fitness than the most deleterious mutation at the same site, as previously reported for the receptor binding domain of the SARS-CoV-2 Spike protein^12^. The larger number of mutations included in our study and the diversity of proteins in which these were present allowed us to identify specific characteristics of such sites. Specifically, such residues were enriched in naturally variable, surface-exposed loops, that had a large fraction of stabilizing mutations. A recent preprint provided an extensive analysis of single, double, and triple codon deletions, as well as different insertions, across the full proteome of the related Enterovirus A71 virus^11^. As observed in our study, the N-terminal regions of 2A and 3A were found to be tolerant to deletions, suggesting the observed trends are generalizable across different enterovirus species.

The majority of DMS studies analyze MFE in a single, highly susceptible cell line. Whether the derived MFE are generalizable to additional cellular environments, and in particular to more restrictive environments, is unclear. To address this, we compared MFE between HeLa-H1 cells and a less-permissive, immune-competent cell line, RPE cells. Interestingly, MFE remained relatively constant across most sites in the proteome, indicating similar selection pressures across the different cellular environments (Figure 5B). However, several capsid sites showed improved fitness in HeLa-H1 cells compared to RPE cells as a result of DAF expression in HeLa-H1 (Figure 5C, D). Hence, entry factors, which frequently show differential expression across different tissues, impart strong selection pressures on viral proteins. It is important to note that any mutations with low fitness in HeLa-H1 but high fitness in RPE cells may have been selected against during the first passage in HeLa-H1 cells and may therefore be missing in our experiments.

Antiviral drug development is a long, complex, and costly process. Unfortunately, the extreme evolutionary capacity of RNA viruses can rapidly render antiviral therapies ineffective^43^. A potential application of DMS can be the identification of protein regions that are intolerant to mutations and can therefore present a higher barrier to the emergence of drug resistance. As a first estimation of whether MFE can be used to inform drug target selection, we incorporated our MFE data into predicted drug pockets across the full proteome. This revealed a highly intolerant site in 3D, where only a single mutation out of all possible mutations across the 34 pocket residues had a neutral MFE, with the remaining mutations being either deleterious or lethal. Such a drug pocket is likely to present a high barrier to the development of resistance due to the high fitness costs incurred by drug-resistant mutations. On the other hand, druggable pockets in the capsid and the mutation-tolerant 2A protein had a notable fraction of neutral mutations suggesting a low barrier to the development of drug resistance. Beyond prioritizing drug targets, MFE data can be used to select mutations that are likely to arise in a given druggable pocket due to low fitness costs. These mutations can then be incorporated into *in silico* drug screening to *a priori* select compounds that will be insensitive to such mutations and to inform medicinal chemistry.

## Acknowledgments

We would like to thank Drs. Rafael Sanjuan and Santiago F. Elena for their helpful suggestions. The computations were performed on the HPC cluster Garnatxa at the Institute for Integrative Systems Biology (I2SysBio), a mixed research center formed by the University of Valencia (UV) and the Spanish National Research Council (CSIC). In addition, the authors would like to acknowledge the Principe Felipe Research Center (CIPF) server for Alphafold2 predictions, which was co-financed by the European Union through the Operativa Program of the European Regional Development Fund (ERDF/FEDER) of the Comunitat Valenciana 2014–2020.

## Author contributions

R.G. acquired funding and conceived the project. B.A-R. and R.G. designed the DMS experiments, analyzed the data, and wrote the manuscript. B.A-R. and S.V-A. performed all experiments. C.T. designed the bioinformatic analysis of deletions.

## RESOURCE AVAILABILITY

### Lead Contact

Further information and requests for resources and reagents should be directed to and will be fulfilled by the lead contact, Ron Geller.

### Materials availability

The materials generated in this work are available from the lead contact with no restrictions.

### Data and code availability

Sequencing data for this project has been deposited in SRA (Bioproject IDs: PRJNA643896, PRJNA779606, PRJNA1013170, PRJNA1033421, see GitHub section B2 for sample details). All scripts used to analyze the data are available on the project’s GitHub (https://github.com/RGellerLab/CVB3_full_proteome_DMS).

## EXPERIMENTAL MODEL AND SUBJECT DETAILS

### Cells, viruses, and reagents

HeLa-H1 (CRL-1958; RRID: CVCL_3334) and RPE (CRL-4000) were obtained from ATCC. Both cell lines were periodically validated to be free of mycoplasma. Cells were cultured in culture media (Dulbecco’s modified Eagle’s medium, Pen-Strep, and L-glutamine) supplemented with 10% or 2% heat-inactivated fetal bovine serum for culturing or infection, respectively. The human codon-optimized T7 polymerase plasmid was obtained from Addgene (#65974) and the CVB3 infectious clones encoding mCherry (CVB3-mCherry) or eGFP (CVB3-eGFP) were previously described^44^. The titer of these reporter viruses was obtained by infecting HeLa-H1 cells with serial dilutions of each virus in 96-well plates and counting the number of fluorescent cells at 8 h post-infection using an Incucyte SX5 Live-Cell Analysis System (Sartorius). All virus experiments were carried out under BSL2 conditions after obtaining approval from the biosafety committees of both the institute and the University of Valencia. All work with genetically modified organisms was approved by the relevant national committees. Antibodies targeting CAR (E-1; sc-373791), CD55/DAF (NaM16-4D3; sc-51733), GAPDH (0411; sc-47724), and goat anti-mouse IgG-HRP (sc-2005) were purchased from Santa Cruz and interferon-β was purchased from Abcam (ab71475).

### Mutant library construction

P1 DMS libraries were obtained as described in^16^. Briefly, the full capsid region of CVB3 was subjected to a PCR-based mutagenesis protocol using degenerate primers, with random bases at the codon-matching positions (NNN), in triplicate. The mutagenized region was cloned into the CVB3 infectious clone and used to generate viral populations. The virus populations were then passaged at low multiplicity of infection (MOI) to create a passage 1 population in which there is a genotype-phenotype link between the capsid proteins of each virus and the genome it carries. These populations were previously characterized in detail^45^.

To generate the P2 and P3 mutagenized regions, a synthetic biology approach was utilized. Briefly, each region was divided into ~250-bp adjacent regions for the mutagenesis and was flanked by complementary sequences to the upstream and downstream regions in the infectious clone. In total, eight and ten such tiles were designed for P2 and P3, respectively. Within each tile, 20 oligonucleotides were ordered for each residue, 19 encoding each of the possible AA mutations and one with the WT codon deleted. Mutated codons were designed to incorporate the maximum number of nucleotide changes versus the wildtype codon and the more frequently utilized codon in the CVB3 ORF was chosen when multiple possibilities were available. In addition, synonymous codons were included at GTG, GAA, AAA, TAC, CAA, and GAG, and stop codons were included at R, L, and S AA as controls. Oligonucleotides were ordered as a single pool for the P2 region and as two pools for the P3 region from Twist Biosciences (See GitHub section A2 for codon choices and full tile sequences). Each tile region was then amplified by PCR using primers matching the overlap region with KAPA HiFi DNA polymerase for 14-16 cycles. PCR products were purified and size-selected using CleanNGS (CleanNA) beads (ratio of 0.75X) and concentrated using DNA clean and concentrator columns (ZYMO research). Finally, the purified PCR was joined to a complementary PCR of the infectious clone using HiFi DNA assembly (See table S1 for primer sequences). In total, three libraries were generated for each tile. Due to the limited quantity of synthetic oligonucleotides, the third replicate of the P2 libraries was amplified by PCR as previously described and cloned in a pJET plasmid for downstream amplification. PJET-cloned tiles were amplified for 14 cycles by KAPA HiFi, size-selected using E-Gel SizeSelect II Gels (Thermo Fisher Scientific), and purified using CleanNGS (CleanNA) beads (1X). Tiles were cloned into the CVB3 infectious clone using HiFi DNA assembly as previously described^16^.

The assembled plasmid reactions were purified using a Zymo DNA Clean and Concentrator-5 kit (Zymo Research) and electroporated into bacteria as previously described^16^. Cells were then grown overnight in a 10 mL liquid culture at 37°C and DNA purified using the NZYMiniprep kit (NZYtech). Transformation efficiency was estimated by plating serial dilutions of the transformation on agar plates. In total, 6- 18*10^5^ transformants were obtained for each line. Viral genomic RNA was then in vitro transcribed and electroporated into HeLa-H1 cells as previously described^16^. Electroporated cells were then pooled and cultured for 9 hr to produce the passage 0 virus (P0). Following three freeze–thaw cycles, cell debris were removed by centrifugation, and viral titer was obtained by plaque assay. Two additional rounds of passaging were performed in HeLa-H1 cells using 10^6^ PFUs (MOI of 0.1) and allowing infection to continue for a single cycle (8h). All infections in HeLa-H1 produced > 1.30×10^7^ PFU in P0 and > 2.45×10^7^ PFU in P1 as judged by plaque assay. For RPE infection, the passage 1 virus populations produced above were used to infect 5×10^6^ cells using 1×10^7^ PFU for 8h. The viral titer of passage 2 for all the RPE infections was >5×10^6^ PFU.

### Duplex sequencing

For sequencing, viral RNA extracted as indicated above was reverse transcribed with the high-fidelity OneScript Plus Reverse Transcriptase (Applied Biological Materials) using 8 μL of RNA and the primers P2_RT_5195 (AATGAAAGCCCGACTGACATGTTT) or CV_7414_R (TTTTTTTTTTTTTTCCGCAC) for P2 or P3 regions, respectively. The number of copies was quantified via qPCR using PowerUp SYBR Green Master Mix (ThermoFisher Scientific) in a 10 μL total reaction, using 1 μL of the template and the primers CVB3P1_qPCR_Fwd (GGAAGCACGGGTCCAATAAA) and CVB3P1_qPCR_Rev (CAGAGTCTAGGTGGTCTAGGTATC) for the P2 region or CVB3P2_qPCR_F (ATGGCTGCCCTAGAAGAGA) and CVB3P2_qPCR_R (CTGACACGGTTGGAGCATTA) for the P3 region. A standard curve was generated using the CVB3 infectious clone plasmid. The full region of interest was then amplified from >10^6^ copies of cDNA using Phusion polymerase (ThermoFisher Scientific) and the primers CVB3_P2_seq_F (AACGTGAACTTCCAACCCAGC) and CVB3_P2_seq_R (CGATTTGAGCAGGTCCGCAAT) to amplify the P2 region or the primers CVB3_P3_seq_F (TCTTGTGTGTGGGAAGGCTATACAAT) and CVB3_P3_seq_R (ACCCCTACTGTACCGTTATCTGGTT) to amplify the P3 region. Duplex sequencing libraries were prepared as previously described^16^ with some modifications. New adaptors were developed that reduce random tag length to 8 instead of 12 and enable dual indices to eliminate index hopping and a synthetic fragment was used to calibrate the amount of DNA used for generating the libraries (see supplementary protocol in GitHub section A1). Samples were sequenced on an Illumina Novaseq 6000 sequencer. The resulting files were analyzed as previously described^45^ using a modified version of the duplex sequencing pipeline^46^, except that a modification was introduced into the VirVarSeq script^47^ to enable the identification of codon deletions (full analysis scripts including the modified VirVarSeq script are available on GitHub, section B2) and the 2021 version of the duplex UnifiedConsensusMaker.py script was used (https://github.com/Kennedy-Lab-UW/Duplex-Seq-Pipeline; minimum family size of 2 and 70% agreement within family required to call a mutation). With this, the counts of each codon at each position were obtained (codon tables, available on GitHub, section A3).

### Calculation and visualization of MFE

As before^16^, all single mutations in codons were omitted from the analysis to increase the signal-to-noise ratio in the case of P1. For P2 and P3, codon tables were filtered to keep only those codons where mutations were introduced. MFE were calculated following the general procedure described in dms_tools2 for the ratio method ^48^ (https://jbloomlab.github.io/dms_tools2/index.html) using custom scripts (available on GitHub, section B3). Briefly, for each site, the sum of all codons giving rise to a particular AA mutation was divided by that of the WT AA to obtain the relative enrichment of each mutation at each site. To obtain the MFE, the relative enrichment for a given mutation in the viral populations was divided by its relative enrichment in the mutagenized libraries. To avoid zeros in the numerator, a coverage-scaled pseudocount of 1 was added to the count of each mutation. Additionally, all mutations which were not observed at least 5 times in the mutagenized libraries were omitted from analysis and mutations observed at least 5 times in the mutagenized libraries and 0 times in the viral populations were defined as lethal. Finally, MFE from replicate lines were then averaged to obtain the average MFE of each mutation, as long as at least 2 MFE values were observed. To normalize MFE between regions, MFE belonging to the overlap regions were used to fit a linear model with the lm function in R. These were then applied to all MFE from the corresponding region (P1 or P3; see GitHub section B4 for data and scripts). Overlap-normalized MFE were used to obtain a linear model for DMS-derived MFE relative to experimental fitness measurements following outlier removal with the Cook’s Distance method (threshold for outliers: Cook’s distance > 4/n; where n is the total number of observations). This transformation was applied to the full dataset (see GitHub section B4 for data and scripts). For downstream analyses lethal mutations were assigned the minimum MFE value observed in the full proteome. All MFE plots in the main figures represent the mean MFE for each mutation at each site across replicates. Line plots represent the mean MFE per site.

To calculate differential selection between HeLa-H1 and RPE cells, MFE were obtained as indicated above. In this case, mutations that were not observed at least 5 times in one of the two viral populations were omitted from the analysis. MFE from HeLa-H1 cells were divided by those from RPE cells to obtain a mutation differential selection score, with values >1 and <1 indicating improved fitness in HeLa-H1 or RPE cells. To obtain the site differential selection scores, all mutations having a mutation differential selection score >1 or <1 were summed to define the site differential selection score for HeLa-H1 or RPE cells, respectively. Scripts to perform the differential selection analysis are available on GitHub (section B5).

### Competition assays to assess the fitness of virus mutants

CVB3 mutants were generated by site-directed mutagenesis using the mCherry-CVB3 fluorescent infectious clone mentioned above as previously described^45^ and utilized in competition assays as previously described^45^. Briefly, all mutants were cloned into the CVB3-mCherry infectious clone and recovered in HEK293T cells. These mutants or the WT mCherry reporter virus were mixed at a 1:1 ratio with a GFP-expressing CVB3 reference virus and used to infect HeLa-H1 cells at an MOI of 0.001 in a 24-well plate. The number of cells infected with each virus (GFP^+^ or mCherry^+^) was determined at both 8hpi (initial infection ratio) and 20hpi (following ~2 rounds of replication) using a live cell microscope (Incucyte SX5; Sartorius). The relative fitness of each mutant or the WT mCherry virus was calculated relative to the GFP-expressing reference virus using the formula (mCherry^+^20h/GFP^+^20h)/(mCherry^+^8h/GFP^+^8h). Fitness values were then standardized relative to that of the WT virus. The results of the competition assay are available on GitHub (section A4).

### Cell-susceptibility, interferon sensitivity, and receptor expression analyses

For calculation of cell susceptibility, HeLa-H1, and RPE cells were infected with identical quantities of an mCherry-expressing CVB3, and virus titer was determined by examination of fluorescence at 8 hours post-infection on a live-cell microscope (Incucyte SX5; Sartorius). For evaluation of type I interferon sensitivity, cells were mock treated or treated with 20 U/mL interferon-β (ab71475; Abcam) for 24 hours. Subsequently, the cells were infected with CVB3-mCherry. The number of infected cells was quantified by examining mCherry^+^ cells using a live cell microscope (Incucyte SX5; Sartorius). For analysis of receptor expression, 5×10^6^ cells were lysed using 0.5 mL of lysis buffer (50mM HEPES pH 7.2, 1% NP-40) containing protease inhibitor cocktail (78430; Thermo Fisher Scientific) on ice for 5 minutes. Nuclei were then separated by centrifugation (10,000 x g for 5 minutes) at 4°C. The lysates were mixed with Laemmli buffer containing β-mercaptoethanol (1610747; Bio-Rad), denatured at 95°C for 5 minutes, separated on a 4-20% gradient precast gel (4561095; Bio-Rad), and transferred to a PVDF membrane. Subsequently, the membrane was blocked with TBS-T containing 3% of BSA, incubated with the relevant primary antibodies for 1 hour (anti-DAF: SC-51733, anti-CAR: SC-373791, anti-GAPDH: SC-47724; Santa Cruz Biotechnology), washed three times with TBS-T and incubated with the secondary antibody for 1 hour (anti-mouse-HRP: SC-2005; Santa Cruz Biotechnology). Finally, the membrane was washed three times, and the chemiluminescent substrate was added (34577; Thermo Fisher Scientific). Quantification of protein bands was done with ImageJ.

### Bioinformatic analyses

For examination of sequence variability in enterovirus B alignments, all available sequences matching the criteria “taxon id 138949, minimum size 2180, 0 ambiguous characters, host homo sapiens, and exclude lab host”, were downloaded from NCBI virus on February 8^th^, 2024. Sequences were then filtered in R to remove those with non-ATGC characters (n = 1,042 remaining), translated to AA sequences using the translate function from the Biostrings package (Version 2.70.1) and aligned to the CVB3 Nancy ORF using the AlignSeqs function of the DECIPHER package (version 2.30.0). All positions with gaps in CVB3 were removed from the alignment (see GitHub section A6 for alignment), and AA counts at each position were summed using the ConsensusMatrix function of Biostrings. Finally, Shannon’s entropy of AA was calculated using the formula −sum(log(p[p>0]/sum(p)) * (p[p>0]/sum(p))), where p is the frequency of a given AA. Phydms^17^ was used to evaluate if the incorporation of MFE into phylogenetic models could improve model fit compared to standard models. For this, all available CVB3 sequences were downloaded on February 8^th^, 2024 using the abovementioned criteria but with taxon id 12072. Sequences were aligned and processed as indicated above for entropy measurements and split into P1, P2, or P3 regions. MFE values were transformed into site preferences by first setting all missing values to the average MFE at a particular site and normalized by dividing by the sum of all MFE values at each site. Phydms was then run using the default setting following preparation of the sequence alignment with the phydms_prepalignment function (see GitHub A6 for subalignments and AA site preferences).

The CVB3 capsid structure was based on the PDB:4GB3, but modified to match the Nancy sequence as previously reported^16^. Structure prediction for all remaining CVB3 proteins was performed using Alphafold^49^ (version 2.0.1), with parameters max_template_date=2020-05-14 and uniclust30_2018_08. The multimer version of Alphafold^50^ was similarly run for all multimeric proteins using the relevant number of subunits. For 2B, the dimeric prediction yielded more similar results to the expected structure defined by NMR^30^ than the tetrameric prediction and was used. Protein stability calculations were obtained from a prior analysis for the capsid proteins^16^, or performed on the Alphafold2 predicted structures using MutateX^51^. To calculate the Stability index, the number of neutral or stabilizing mutations (DDG<1) at each residue was subtracted from the number of residues. RSA and secondary structure were obtained from PSAIA^52^ and STRIDE^53^, respectively, using the Alphafold2 predicted protein structures. Residues with RSA>0.25 were classified as surface exposed. Interface residues were obtained using the interfaceResidues script in PyMOL (cutoff of interface accessible surface area = 0.75 Å²). Identification of druggable pockets was performed using the Schrödinger software (Version 13.8). Proteins were first subjected to the *protein preparation workflow* in Maestro using the default settings and then SiteMap was used to define druggable pockets using the default settings, with sites having a SiteMap score >0.8 considered as druggable.

### Statistical analyses

All statistical tests were performed in R (version 4.3.2) and were two-tailed. Multiple testing corrections were performed using the FDR method.

## REFERENCES

1. I, W.Y., Darius, K., Jaime, I., Adriana, L.-S., H, K.J., Mart, K., V, D.V., and V, K.E. (2018). Origins and Evolution of the Global RNA Virome. mBio 9, 10.1128/mbio.02329-18. 10.1128/mbio.02329-18.

2. Peck, K.M., and Lauring, A.S. (2018). Complexities of Viral Mutation Rates. J Virol 92, 1–8. 10.1128/jvi.01031-17.

3. Sanjuan, R., and Domingo-Calap, P. (2016). Mechanisms of viral mutation. Cellular and Molecular Life Sciences 73, 4433–4448. 10.1007/s00018-016-2299-6.

4. Sanjuán, R., Nebot, M.R., Chirico, N., Mansky, L.M., and Belshaw, R. (2010). Viral mutation rates. J Virol 84, 9733–9748. 10.1128/JVI.00694-10.

5. Rangel, M.A., Dolan, P.T., Taguwa, S., Xiao, Y., Andino, R., and Frydman, J. (2023). High-resolution mapping reveals the mechanism and contribution of genome insertions and deletions to RNA virus evolution. Proc Natl Acad Sci U S A 120. 10.1073/pnas.2304667120.

6. Geller, R., Estada, Ú., Peris, J.B., Andreu, I., Bou, J.-V., Garijo, R., Cuevas, J.M., Sabariegos, R., Mas, A., and Sanjuán, R. (2016). Highly heterogeneous mutation rates in the hepatitis C virus genome. Nat Microbiol 1, 16045. 10.1038/nmicrobiol.2016.45.

7. Lorenzo-Redondo, R., de Sant’Anna Carvalho, A.M., Hultquist, J.F., and Ozer, E.A. (2023). SARS-CoV-2 genomics and impact on clinical care for COVID-19. Journal of Antimicrobial Chemotherapy 78, ii25–ii36. 10.1093/jac/dkad309.

8. Raman, H.S.A., Tan, S., August, J.T., and Khan, A.M. (2020). Dynamics of Influenza A (H5N1) virus protein sequence diversity. PeerJ 2020. 10.7717/peerj.7954.

9. Gabrielaite, M., Bennedbæk, M., Rasmussen, M.S., Kan, V., Furrer, H., Flisiak, R., Losso, M., Lundgren, J.D., and Marvig, R.L. (2023). Deep-sequencing of viral genomes from a large and diverse cohort of treatment-naive HIV-infected persons shows associations between intrahost genetic diversity and viral load. PLoS Comput Biol 19. 10.1371/journal.pcbi.1010756.

10. Acevedo, A., Brodsky, L., and Andino, R. (2014). Mutational and fitness landscapes of an RNA virus revealed through population sequencing. Nature 505, 686–690. 10.1038/nature12861.

11. Bakhache, W., Walker, O., McCormick, L., and Dolan, P. (2024). Uncovering Structural Plasticity of Enterovirus A through Deep Insertional and Deletional Scanning. PREPRINT. 10.21203/rs.3.rs-3835307/v1.

12. Taylor, A.L., and Starr, T.N. (2023). Deep mutational scans of XBB.1.5 and BQ.1.1 reveal ongoing epistatic drift during SARS-CoV-2 evolution. PLoS Pathog 19, e1011901-.

13. Ogden, P.J., Kelsic, E.D., Sinai, S., and Church, G.M. (2019). Comprehensive AAV capsid fitness landscape reveals a viral gene and enables machine-guided design. Science 366, 1139–1143. 10.1126/science.aaw2900.

14. Racaniello, V.R. (2013). Picornaviridae: The Viruses and Their Replication. In Fields Virology, M. D. Knipe and M. P. Howley, eds. (Philadelphia : Wolters Kluwer Health/Lippincott Williams & Wilkins, c2013.), pp. 453–489.

15. Burton, T.D., and Eyre, N.S. (2021). Applications of Deep Mutational Scanning in Virology. Viruses 13. 10.3390/v13061020.

16. Mattenberger, F., Latorre, V., Tirosh, O., Stern, A., and Geller, R. (2021). Globally defining the effects of mutations in a picornavirus capsid. Elife 10, e64256. 10.7554/eLife.64256.

17. Hilton, S.K., Doud, M.B., and Bloom, J.D. (2017). Phydms: Software for phylogenetic analyses informed by deep mutational scanning. PeerJ. 10.7717/peerj.3657.

18. Jiang, P., Liu, Y., Paul, A. V., Wimmer, E., Ma, H.-C., Paul, A. V., and Wimmer, E. (2014). Picornavirus Morphogenesis. Microbiology and Molecular Biology Reviews 78, 418–437. 10.1128/MMBR.00012-14.

19. Álvarez-Rodríguez, B., Buceta, J., and Geller, R. (2023). Comprehensive profiling of neutralizing polyclonal sera targeting coxsackievirus B3. Nat Commun 14. 10.1038/s41467-023-42144-2.

20. Mondal, S., Sarvari, G., and Boehr, D.D. (2023). Picornavirus 3C Proteins Intervene in Host Cell Processes through Proteolysis and Interactions with RNA. Viruses 15, 2413. 10.3390/v15122413.

21. Yang, X., Cheng, A., Wang, M., Jia, R., Sun, K., Pan, K., Yang, Q., Wu, Y., Zhu, D., Chen, S., et al. (2017). Structures and corresponding functions of five types of picornaviral 2A proteins. Preprint at Frontiers Media S.A., 10.3389/fmicb.2017.01373 10.3389/fmicb.2017.01373.

22. Liu, Y., Li, J., and Zhang, Y. (2023). Update on enteroviral protease 2A: Structure, function, and host factor interaction. Preprint at Elsevier B.V., 10.1016/j.bsheal.2023.09.001 10.1016/j.bsheal.2023.09.001.

23. Zhang, X., Paget, M., Wang, C., Zhu, Z., and Zheng, H. (2020). Innate immune evasion by picornaviruses. Eur J Immunol 50, 1268–1282. 10.1002/eji.202048785.

24. Ypma-Wong, M.F., Dewalt, P.G., Johnson, V.H., Lamb, J.G., and Semler, B.L. (1988). Protein 3CD is the major poliovirus proteinase responsible for cleavage of the p1 capsid precursor. Virology 166, 265–270. 10.1016/0042-6822(88)90172-9.

25. Geller, R., Pechmann, S., Acevedo, A., Andino, R., and Frydman, J. (2018). Hsp90 shapes protein and RNA evolution to balance trade-offs between protein stability and aggregation. Nat Commun 9, 1781. 10.1038/s41467-018-04203-x.

26. Wolff, G., Melia, C.E., Snijder, E.J., and Bárcena, M. (2020). Double-Membrane Vesicles as Platforms for Viral Replication. Preprint at Elsevier Ltd, 10.1016/j.tim.2020.05.009 10.1016/j.tim.2020.05.009.

27. Suhy, D.A., Giddings, T.H., and Kirkegaard, K. (2000). Remodeling the Endoplasmic Reticulum by Poliovirus Infection and by Individual Viral Proteins: an Autophagy-Like Origin for Virus-Induced Vesicles. J Virol 74. 10.1128/jvi.74.19.8953-8965.2000.

28. Jackson, T., and Belsham, G.J. (2021). Review picornaviruses: A view from 3a. Viruses 13, 1–42. 10.3390/v13030456.

29. Wang, S.H., Wang, K., Zhao, K., Hua, S.C., and Du, J. (2020). The Structure, Function, and Mechanisms of Action of Enterovirus Non-structural Protein 2C. Preprint at Frontiers Media S.A., 10.3389/fmicb.2020.615965 10.3389/fmicb.2020.615965.

30. Li, Z., Zou, Z., Jiang, Z., Huang, X., and Liu, Q. (2019). Biological function and application of picornaviral 2B protein: A new target for antiviral drug development. Viruses 11, 1–16. 10.3390/v11060510.

31. Xia, X., Cheng, A., Wang, M., Ou, X., Sun, D., Mao, S., Huang, J., Yang, Q., Wu, Y., Chen, S., et al. (2022). Functions of Viroporins in the Viral Life Cycle and Their Regulation of Host Cell Responses. Preprint at Frontiers Media S.A., 10.3389/fimmu.2022.890549 10.3389/fimmu.2022.890549.

32. Li, Z., Zou, Z., Jiang, Z., Huang, X., and Liu, Q. (2019). Biological function and application of picornaviral 2B protein: A new target for antiviral drug development. Preprint at MDPI AG, 10.3390/v11060510 10.3390/v11060510.

33. Wang, S.H., Wang, K., Zhao, K., Hua, S.C., and Du, J. (2020). The Structure, Function, and Mechanisms of Action of Enterovirus Non-structural Protein 2C. Preprint at Frontiers Media S.A., 10.3389/fmicb.2020.615965 10.3389/fmicb.2020.615965.

34. Hurdiss, D.L., El Kazzi, P., Bauer, L., Papageorgiou, N., Ferron, F.P., Donselaar, T., Van Vliet, A.L.W., Shamorkina, T.M., Snijder, J., Canard, B., et al. (2022). Fluoxetine targets an allosteric site in the enterovirus 2C AAA+ ATPase and stabilizes a ring-shaped hexameric complex. Sci Adv 8. 10.1126/sciadv.abj7615.

35. Melia, C.E., van der Schaar, H.M., Lyoo, H., Limpens, R.W.A.L., Feng, Q., Wahedi, M., Overheul, G.J., van Rij, R.P., Snijder, E.J., Koster, A.J., et al. (2017). Escaping Host Factor PI4KB Inhibition: Enterovirus Genomic RNA Replication in the Absence of Replication Organelles. Cell Rep 21, 587–599. 10.1016/j.celrep.2017.09.068.

36. Peersen, O.B. (2017). Picornaviral polymerase structure, function, and fidelity modulation. Preprint at Elsevier B.V., 10.1016/j.virusres.2017.01.026 10.1016/j.virusres.2017.01.026.

37. Thompson, A.A., and Peersen, O.B. (2004). Structural basis for proteolysis-dependent activation of the poliovirus RNA-dependent RNA polymerase. EMBO Journal 23, 3462–3471. 10.1038/sj.emboj.7600357.

38. Xiao, Y., Dolan, P.T., Goldstein, E.F., Li, M., Farkov, M., Brodsky, L., and Andino, R. (2017). Poliovirus intrahost evolution is required to overcome tissue-specific innate immune responses. Nat Commun 8, 375. 10.1038/s41467-017-00354-5.

39. Bergelson, J.M., Cunningham, J.A., Droguett, G., Kurt-Jones, E.A., Krithivas, A., Hong, J.S., Horwitz, M.S., Crowell, R.L., and Finberg, R.W. (1997). Isolation of a common receptor for coxsackie B viruses and adenoviruses 2 and 5. Science (1979) 275. 10.1126/science.275.5304.1320.

40. Shafren, D.R., Bates, R.C., Agrez, M. V, Herd, R.L., Burns, G.F., and Barry, R.D. (1995). Coxsackieviruses B1, B3, and B5 use decay accelerating factor as a receptor for cell attachment. J Virol 69. 10.1128/jvi.69.6.3873-3877.1995.

41. Fitzsimmons, W.J., Woods, R.J., McCrone, J.T., Woodman, A., Arnold, J.J., Yennawar, M., Evans, R., Cameron, C.E., and Lauring, A.S. (2018). A speed–fidelity trade-off determines the mutation rate and virulence of an RNA virus. PLoS Biol 16, 1–20. 10.1371/journal.pbio.2006459.

42. Tokuriki, N., Oldfield, C.J., Uversky, V.N., Berezovsky, I.N., and Tawfik, D.S. (2009). Do viral proteins possess unique biophysical features? Trends Biochem Sci 34, 53–59. 10.1016/j.tibs.2008.10.009.

43. Ma, Y., Frutos-Beltrán, E., Kang, D., Pannecouque, C., De Clercq, E., Menéndez-Arias, L., Liu, X., and Zhan, P. (2021). Medicinal chemistry strategies for discovering antivirals effective against drug-resistant viruses. Preprint, 10.1039/d0cs01084g 10.1039/d0cs01084g.

44. Bou, J.-V., Geller, R., and Sanjuán, R. (2019). Membrane-Associated Enteroviruses Undergo Intercellular Transmission as Pools of Sibling Viral Genomes. Cell Rep 29, 714–723.e4. 10.1016/j.celrep.2019.09.014.

45. Álvarez-Rodríguez, B., Buceta, J., and Geller, R. (2023). Comprehensive profiling of neutralizing polyclonal sera targeting coxsackievirus B3. Nat Commun 14, 6417. 10.1038/s41467-023-42144-2.

46. Schmitt, M.W., Kennedy, S.R., Salk, J.J., Fox, E.J., Hiatt, J.B., and Loeb, L.A. (2012). Detection of ultra-rare mutations by next-generation sequencing. Proceedings of the National Academy of Sciences 109, 14508–14513. 10.1073/pnas.1208715109.

47. Verbist, B.M.P., Thys, K., Reumers, J., Wetzels, Y., Van Der Borght, K., Talloen, W., Aerssens, J., Clement, L., and Thas, O. (2015). VirVarSeq: A low-frequency virus variant detection pipeline for Illumina sequencing using adaptive base-calling accuracy filtering. Bioinformatics 31, 94–101. 10.1093/bioinformatics/btu587.

48. Bloom, J.D. (2015). Software for the analysis and visualization of deep mutational scanning data. BMC Bioinformatics 16, 168. 10.1186/s12859-015-0590-4.

49. Jumper, J., Evans, R., Pritzel, A., Green, T., Figurnov, M., Ronneberger, O., Tunyasuvunakool, K., Bates, R., Žídek, A., Potapenko, A., et al. (2021). Highly accurate protein structure prediction with AlphaFold. Nature 596. 10.1038/s41586-021-03819-2.

50. Evans, R., O’Neill, M., Pritzel, A., Antropova, N., Senior, A., Green, T., Žídek, A., Bates, R., Blackwell, S., Yim, J., et al. (2022). Protein complex prediction with AlphaFold-Multimer. bioRxiv.

51. Tiberti, M., Terkelsen, T., Degn, K., Beltrame, L., Cremers, T.C., da Piedade, I., Di Marco, M., Maiani, E., and Papaleo, E. (2022). MutateX: an automated pipeline for in silico saturation mutagenesis of protein structures and structural ensembles. Brief Bioinform 23. 10.1093/bib/bbac074.

52. Mihel, J., Šikić, M., Tomić, S., Jeren, B., and Vlahoviček, K. (2008). PSAIA – Protein Structure and Interaction Analyzer. BMC Struct Biol 8, 21. 10.1186/1472-6807-8-21.

53. Heinig, M., and Frishman, D. (2004). STRIDE: A web server for secondary structure assignment from known atomic coordinates of proteins. Nucleic Acids Res 32. 10.1093/nar/gkh429.

