## Supplementary figures for "Mapping the mutational landscape of a full viral proteome reveals distinct profiles of mutation tolerability"

### Supplementary material

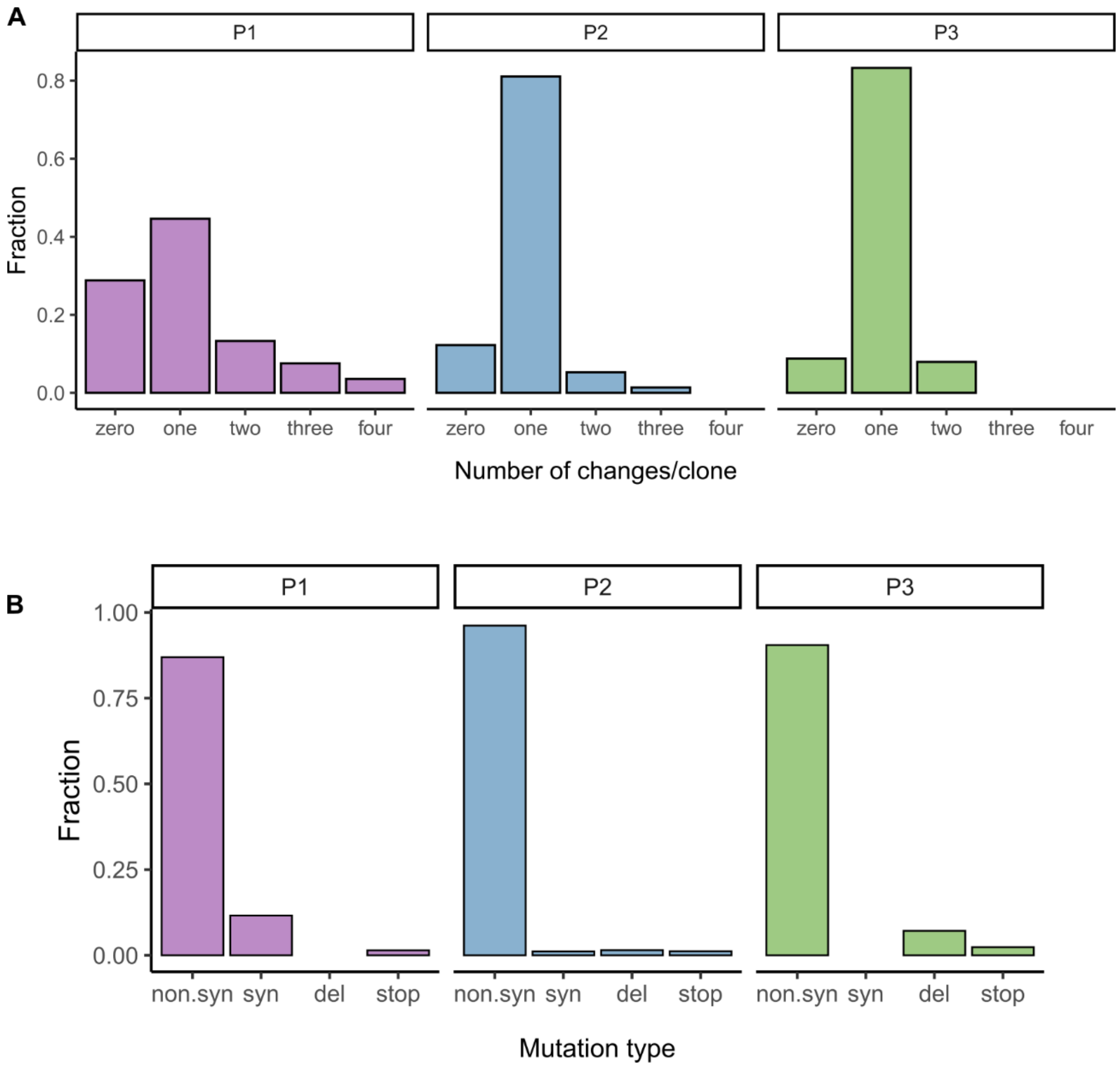


**Figure S1. Related to Figure 1. Characterization of the DMS libraries by Sanger sequencing.** A total of 59, 148, and 92 clones were sequenced for the P1, P2, and P3 region, respectively. The fraction of each number of mutations per clone **(A)** and each type of mutation **(B)** are graphed. Of note, the full capsid region was sequenced for P1, while for P2 and P3, only the corresponding tile region was sequenced.

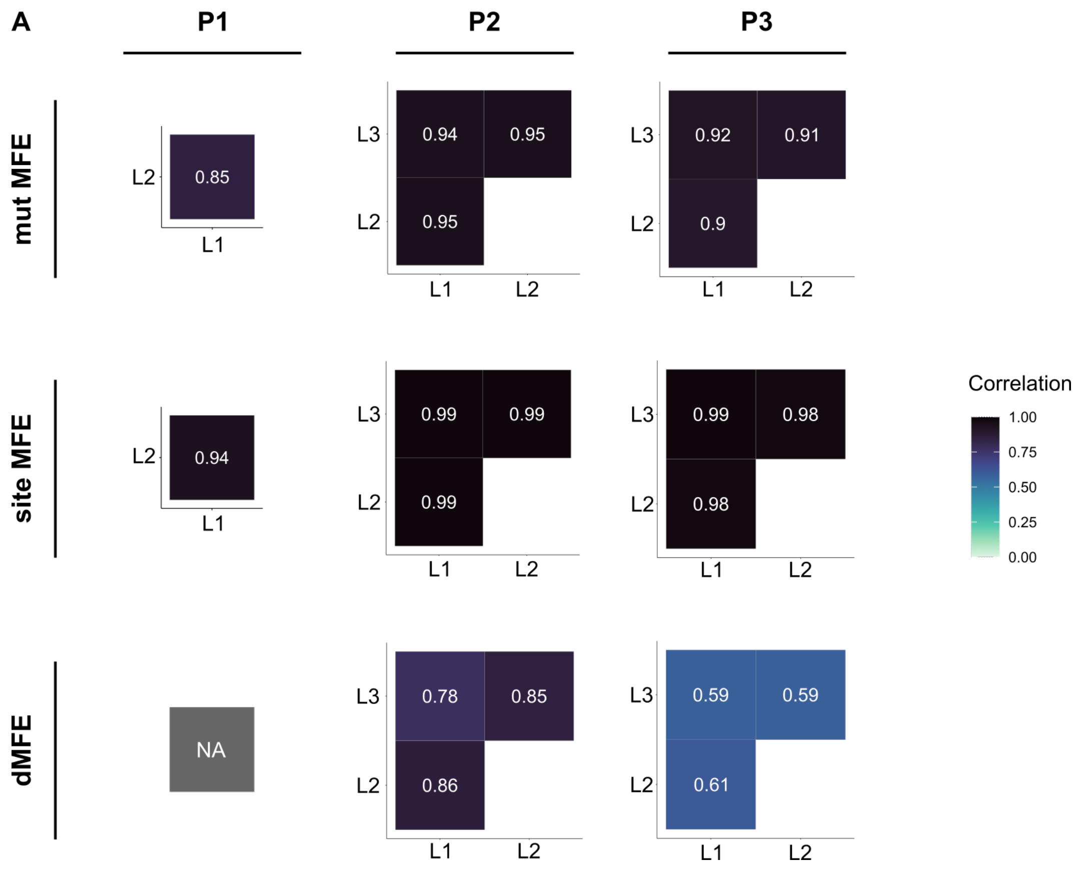


**Figure S2. Related to Figure 1. Correlation of site, mutation and deletion MFEs in Hela-H1 cells between replicates. (A)** Correlations matrices for MFE of mutations (mut MFE), their average per site (site MFE), and deletions (dMFE) for independent replicate lines (L1-L3) for the P1, P2, P3 regions. Of note, for P1, only two replicates were used and deletions were not included in the mutagenesis protocol, precluding their analysis.

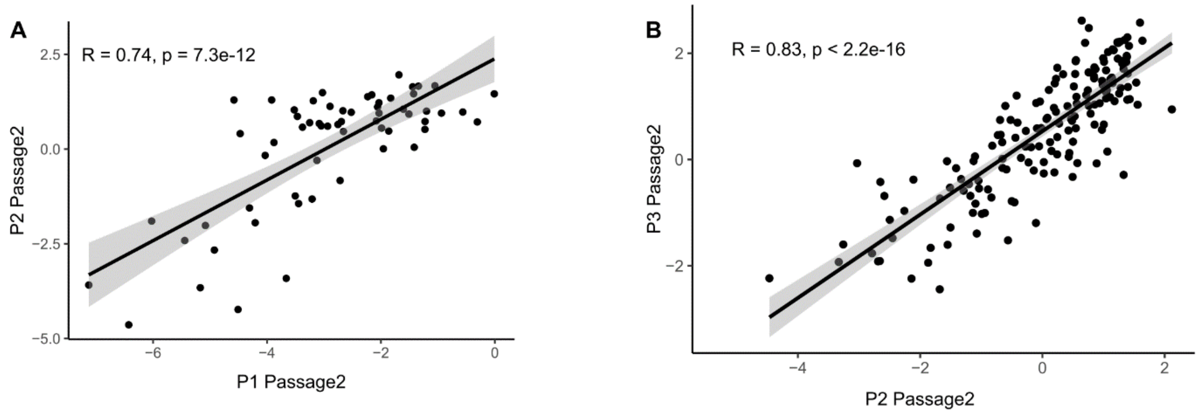


**Figure S3. Related to Figure 2. Linear models used for the normalization of MFE between the different mutagenized regions.** Linear model of MFE present in the overlap between P1 and P2 ($R^2$= 0.54, p = 7.338 x $10^{-12}$) **(A)** or P2 and P3 ($R^2$= 0.69, p < 2.2 x $10^{-16}$) **(B)** used for normalization between regions. The Pearson correlation coefficient and associated p-value are shown.

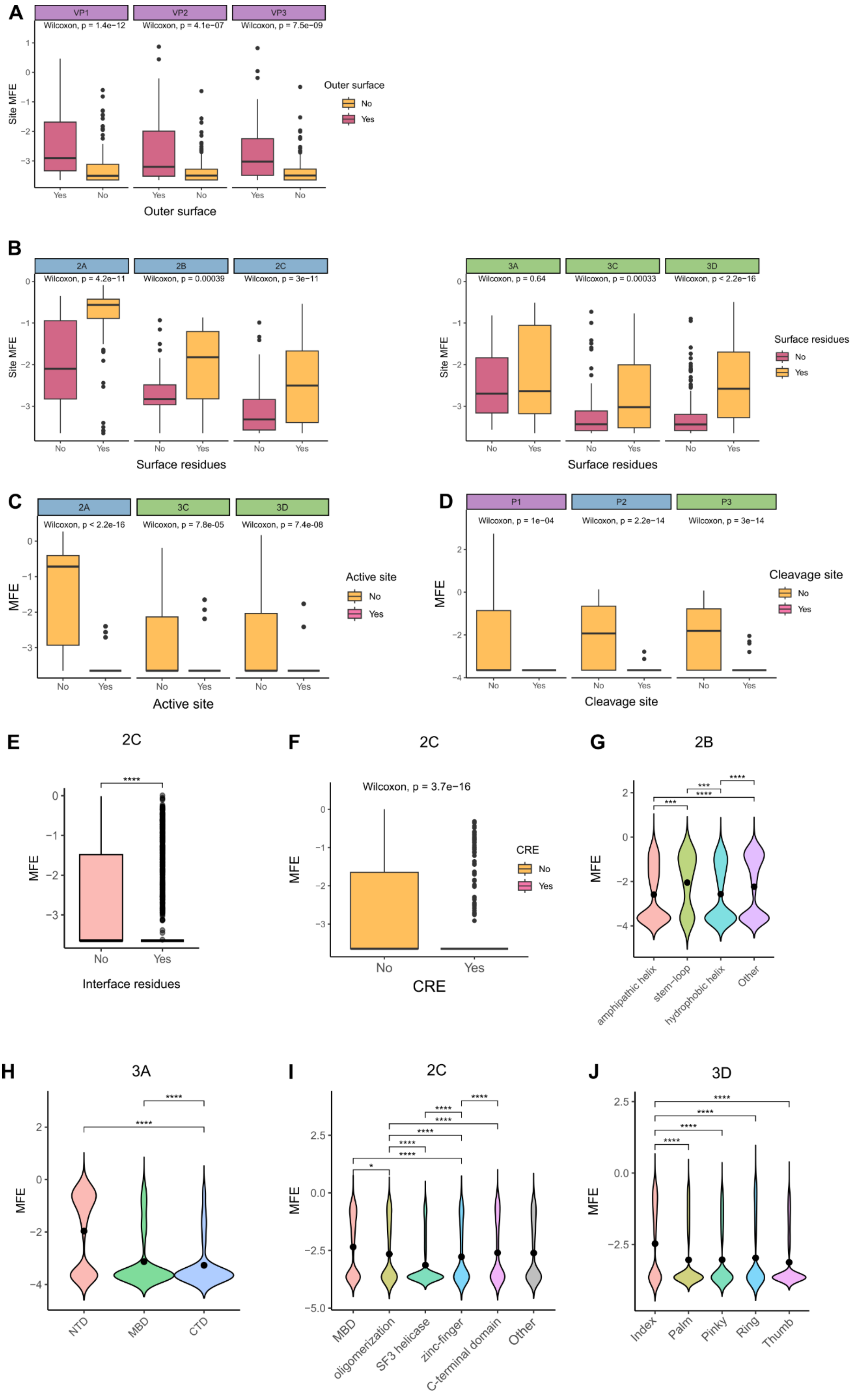
A
VP1
VP2
VP3
Wilcoxon, p = 1.4e−12
Wilcoxon, p = 4.1e−07
Wilcoxon, p = 7.5e−09
Site MFE
Outer surface
No
Yes
B
2A
2B
2C
Wilcoxon, p = 4.2e−11
Wilcoxon, p = 0.00039
Wilcoxon, p = 3e−11
3A
3C
3D
Wilcoxon, p = 0.64
Wilcoxon, p = 0.00033
Wilcoxon, p < 2.2e−16
Surface residues
C
Wilcoxon, p < 2.2e−16
Wilcoxon, p = 7.8e−05
Wilcoxon, p = 7.4e−08
MFE
Active site
D
P1
P2
P3
Wilcoxon, p = 1e−04
Wilcoxon, p = 2.2e−14
Wilcoxon, p = 3e−14
Cleavage site
E
2C
Interface residues
F
Wilcoxon, p = 3.7e−16
CRE
G
2B
amphipathic helix
stem−loop
hydrophobic helix
Other
H
3A
NTD
MBD
CTD
I
oligomerization
SF3 helicase
zinc-finger
C-terminal domain
J
3D
Index
Palm
Pinky
Ring
Thumb

**Figure S4. Related to Figure 3. Protein-specific features of MFE. (A-B)** Distribution of site MFE in outer surface residues versus other residues in the capsid proteins VP1-VP3 **(A)** and in surface exposed residues versus internal residues for the non-structural proteins **(B)**. **(C-D)** Distribution of MFE in the active sites of the 2A and 3C proteases, and the 3D polymerase versus all other residues in each protein **(C)**, and in the 3C protease cleavage site Q residues of each protein versus all other Q residues in that same protein **(D)**. **(E)** Distribution of MFE in interface residues between monomers versus other residues in 2C. **(F)** Distribution of MFE in the CRE element versus other residues in 2C. **(G-J)** Distribution of mutation MFE in the different structural and functional domains of the 2B **(G)**, 3A **(H)**, 2C **(I)**, and 3D **(J)** proteins. ns: $p > 0.05$, *$p < 0.05$, **$p < 0.01$, ***$p < 0.001$, ****$p < 0.0001$ by Mann-Whitney test following multiple test correction.

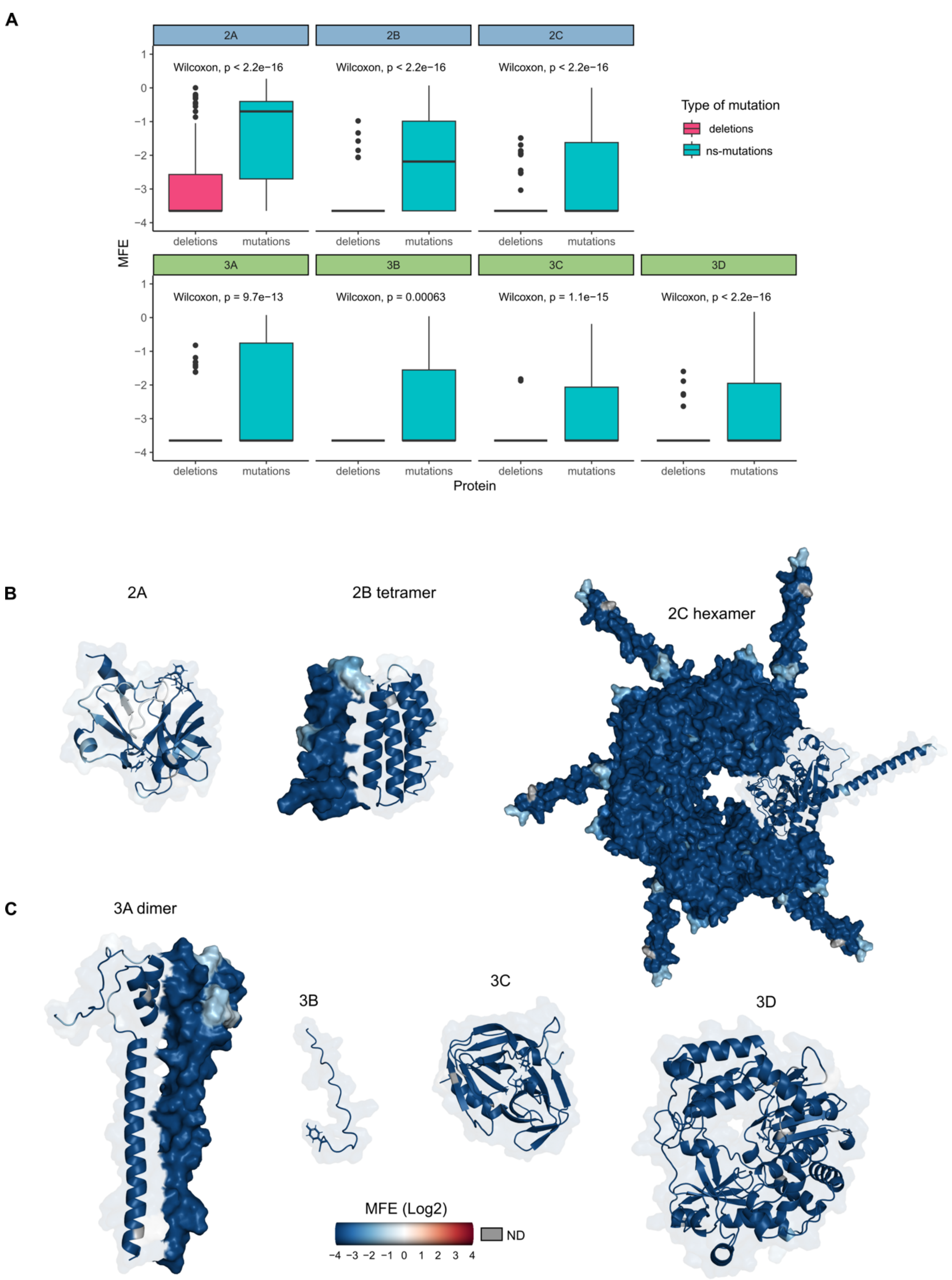


**Figure S5. Related to Figure 4. The fitness effects of single codon deletions across the full CVB3 proteome. (A)** The effects of non-synonymous mutations (ns-mutations) versus deletions across the non-structural proteins. **(B-C)** Mapping of dMFE in P2 **(B)** and P3 **(C)** derived proteins.

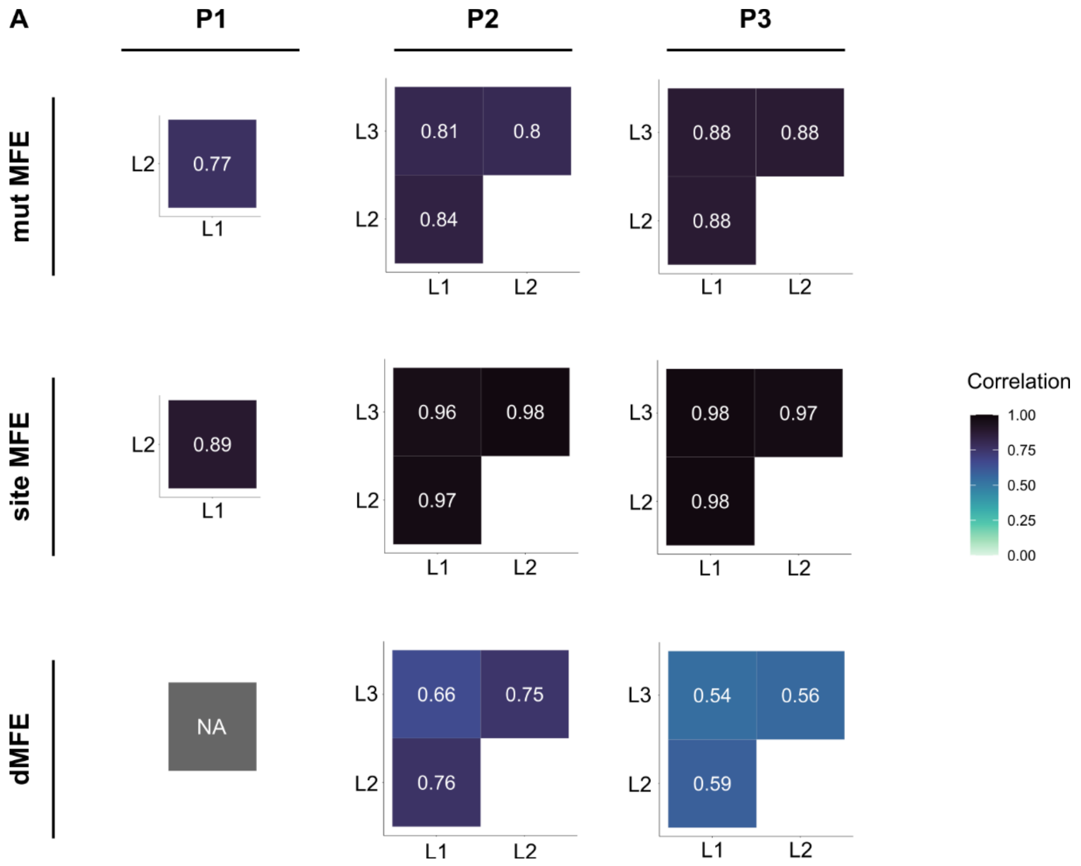


**Figure S6. Related to Figure 5. Correlation of site, mutation and deletion MFE in RPE cells between replicates. (A)** Correlations matrices for MFE of mutations (mut MFE), their average per site (site MFE), and deletions (dMFE) for independent replicate lines (L1-L3) for the P1, P2, P3 regions. Of note, for P1, only two replicates were used and deletions were not included in the mutagenesis protocol, precluding their analysis.

**A** CAR + GAPDH

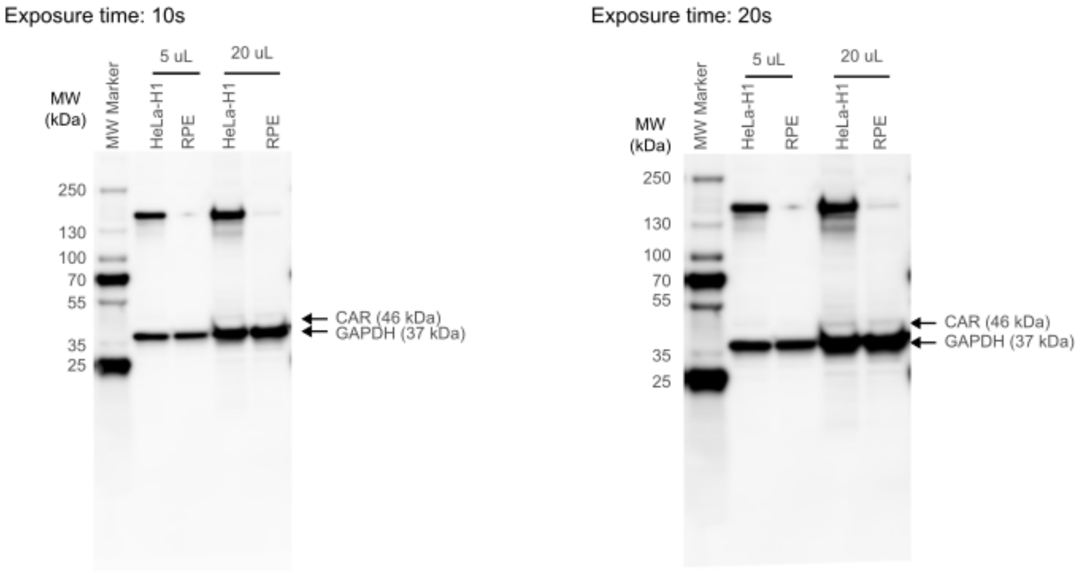


**B** DAF + GAPDH

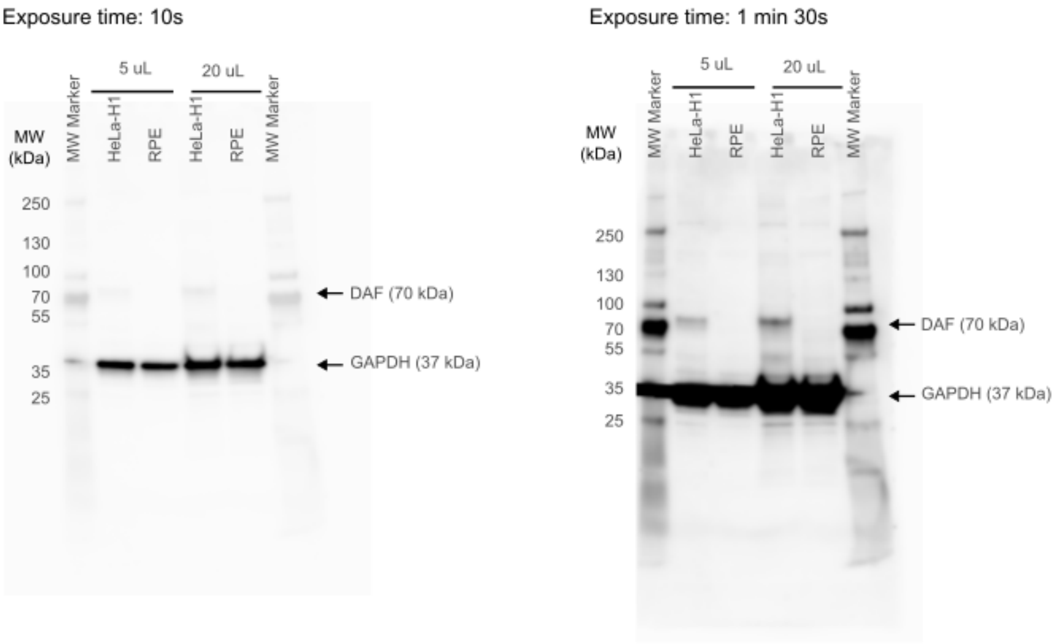


**C**

| Antibody | Exposure time | Band | Hela-H1 | RPE |
|---|---|---|---|---|
| CAR | 1min 30s | CAR | 31935.6 | 34650.8 |
| CAR | 10s | GAPDH | 22981.7 | 25234.8 |
| DAF | 20s | DAF | 13014.0 | 174.6 |
| DAF | 10s | GAPDH | 21351.3 | 19064.1 |

**Figure S7. Related to Figure 5. Analysis of CAR and DAF expression on HeLa-H1 and RPE cells. (A)** The complete membrane is shown for western blots of CAR **(A)** and DAF **(B)** in both cell lines. Arrows indicate bands of the expected size for each protein. A higher molecular weight cross-reactive band is observed for CAR in HeLa-H1. **(C)** Values obtained for the quantification of protein bands. Exposure times of blots used for each protein are indicated.

**Titles of supplementary tables.**

**Table S1. Related to Figure 1. Primers used for the PCR of the synthetic oligonucleotides and the vector to mutagenize regions P2 and P3.**

**Table S2. Related to Figure 1. NGS summary statistics of the DMS plasmid libraries and the virus populations passaged in Hela-H1.**

**Table S3. Related to Figures 2 and 3. Site MFE data, ddg, entropy and annotations for each residue of the CVB3 genome.**

**Table S4. Related to Figures 2, 3 and 4. Mutation MFE data, ddg, entropy and annotations for each possible no-synonymous mutation and deletion of the CVB3 genome.**

**Table S5. Results of the phyDMS analysis for each region of the CVB3 genome.**

**Table S6. Related to Figure 1. NGS summary statistics of the virus populations passaged in RPE.**

**Table S7. Related to Figure 5. Site differential selection values in Hela-H1 and RPE.**

**Table S8. Related to Figure 6. MFE values of predicted druggable pockets.**
